# The bacterial leader peptide peTrpL has a conserved function in antibiotic-dependent posttranscriptional regulation of ribosomal genes

**DOI:** 10.1101/606483

**Authors:** Hendrik Melior, Sandra Maaß, Maximilian Stötzel, Siqi Li, Konrad U. Förstner, Rubina Schütz, Saina Azarderakhsh, Aleksei Shevkoplias, Susanne Barth-Weber, Adithi R. Varadarajan, Kathrin Baumgardt, Zoe Chervontseva, John Ziebuhr, Christian H. Ahrens, Dörte Becher, Elena Evguenieva-Hackenberg

## Abstract

The ribosome-dependent attenuator located upstream of bacterial tryptophan biosynthesis genes harbors a small ORF *trpL* containing tryptophan codons. When tryptophan is available, efficient *trpL* translation causes transcription termination and release of the attenuator RNA rnTrpL. In *Sinorhizobium meliloti*, rnTrpL is a *trans*-acting sRNA. Here, we identified an evolutionary conserved function for the *trpL*-encoded 14-aa leader peptide peTrpL. Upon exposure to tetracycline, the cellular peTrpL levels were increased and rnTrpL was generated independently of tryptophan availability. Both peTrpL and rnTrpL were found to be involved in tetracycline-dependent destabilization of *rplUrpmA* mRNA encoding ribosomal proteins L21 and L27. We provide evidence for redirection of the sRNA rnTrpL from its antibiotic-independent target *trpDC* to *rplUrpmA* by formation of an antibiotic-dependent ribonucleoprotein complex (ARNP). ARNPs comprising peTrpL, rnTrpL, *rplUrpmA* and antisense RNA were also observed for other translation-inhibiting antibiotics, suggesting that bacteria evolved mechanisms to utilize antibiotics for mRNA destabilization.

## Introduction

In bacteria where transcription and translation are coupled, ribosome-dependent transcription attenuation is a conserved regulatory mechanism that relies on small upstream ORFs (uORFs) encoding leader peptides. Despite their widespread occurrence, no functions in *trans* are known for the leader peptides. Here, we assign a function in *trans* to a bacterial leader peptide in the posttranscriptional downregulation of specific ribosomal genes upon exposure to translation-inhibiting antibiotics. We provide evidence that this regulation involves a ribonucleoprotein complex, the formation of which is directly supported by the antibiotics.

Ribosome-dependent transcription attenuators are located upstream of many amino acid (aa) biosynthesis operons in Gram-negative bacteria (Merino et al., 2008; Vitreschak et al., 2004). This posttranscriptional mechanism has been studied most extensively in *Escherichia coli* where the tryptophan (Trp) biosynthesis genes *trpEDCBA* are co-transcribed (Yanofski, 1981). The 5′ mRNA leader harbors the uORF *trpL*, which contains two consecutive Trp codons (codons 10 and 11 of the 14-codon uORF). Under conditions of Trp insufficiency, ribosome pausing at the Trp codons prevents the formation of a transcriptional terminator downstream of the uORF, thus resulting in co-transcription of *trpL* with the structural *trp* genes. In contrast, if enough Trp is available, fast translation of the Trp codons causes transcription termination between *trpL* and *trpE*. However, transcription termination between *trpL* and *trpE* is also caused by ribosome pausing in the first half of *trpL* (Zurawski et al., 1978).

Generally, the released products of ribosome-dependent attenuators, i.e. the small attenuator RNAs and leader peptides, have been considered nonfunctional. Recently, however, we reported that in the soil-dwelling, nitrogen-fixing and Gram-negative plant symbiont *S. meliloti*, the attenuator RNA rnTrpL is a *trans*-acting small RNA (sRNA) (Melior et al., 2019). Like in many other bacteria, the *trp* genes of *S. meliloti* are organized into several operons, only one of which is regulated by attenuation (Merino et al., 2008). The *trp* attenuator of *S. meliloti* is located upstream of *trpE(G)* (Fig. 1A; Bae and Crawford, 1990), which is transcribed separately from *trpDC* and *trpFBA* (Jonston et al., 1978). Under conditions of Trp availability, fast translation of three consecutive Trp codons in *trpL* (codons 9 to 11) results in transcription termination. The released sRNA rnTrpL (110 nt) base pairs with and destabilizes *trpDC* mRNA, thereby posttranscriptionally coordinating the expression of the *trpE(G)* and *trpDC* operons (Melior et al., 2019).

**Figure 1.**
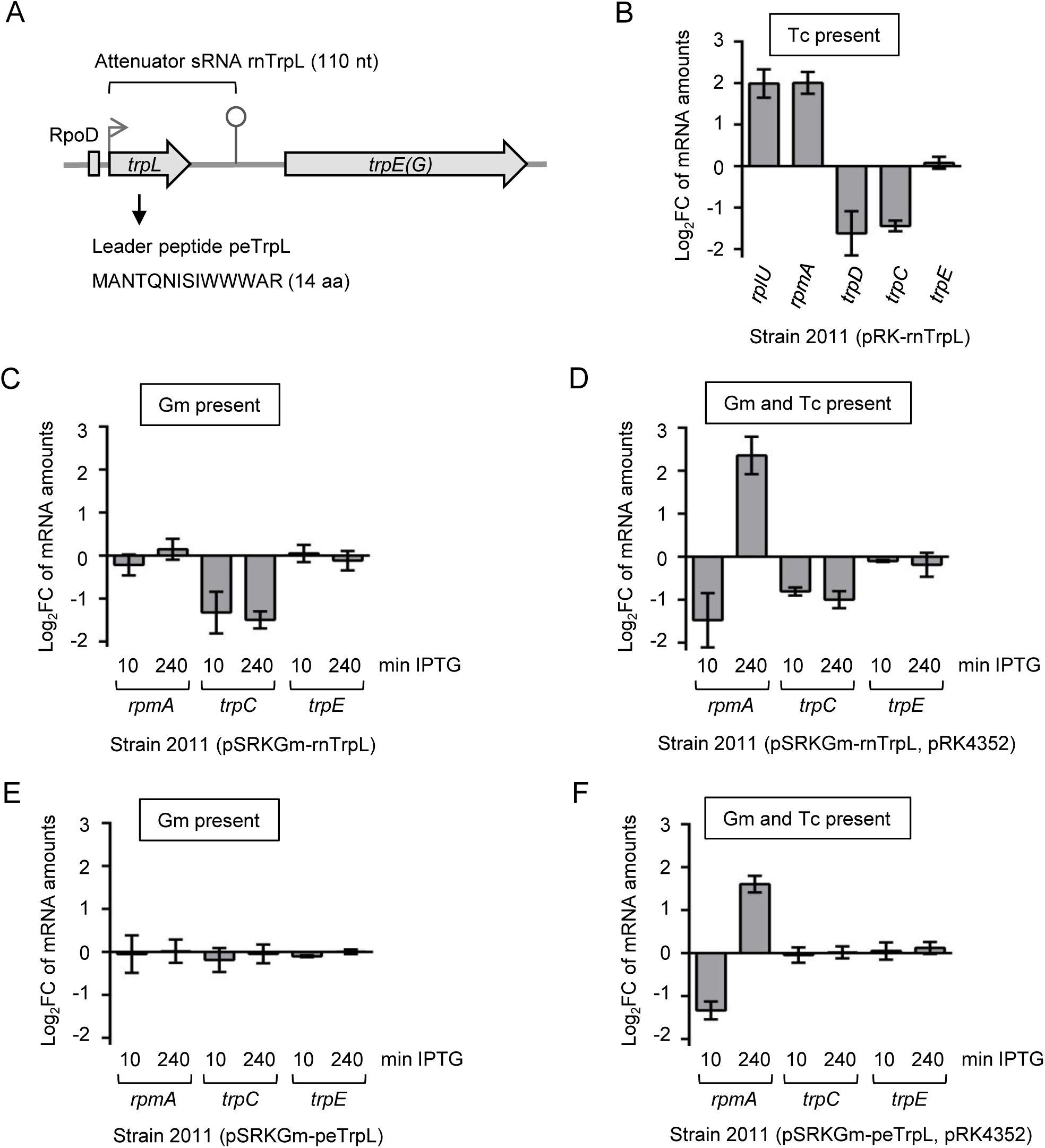
Induced, short-term overproduction of the leader peptide peTrpL decreases the *rplUrpmA* mRNA level in *S. meliloti* 2011. **A)** Schematic representation of the *trpLE(G)* locus. The *trpL* and *trpE(G)* ORFs (gray arrows), a RpoD-like promoter (rectangle), the transcription start site (flexed arrow) and the transcription terminator (hairpin) are depicted (according to Bae and Crawford, 1990). **B)** to **F)** Log_2_ fold changes (log_2_FC) in mRNA levels determined by qRT-PCR. Each graph contains data from three independent experiments, each performed in duplicates (shown are means and standard deviations). In each panel, used overexpression strain and presence of selecting antibiotics in the growth media are indicated. **B)** Changes in the levels of the indicated mRNAs in the constitutively overexpressing strain 2011 (pRK-rnTrpL), in comparison to the EVC 2011 (pRK4352). **C)** and **D)** Changes in the levels of the indicated mRNAs upon induction of rnTrpL overproduction. The sRNA rnTrpL encodes the peptide peTrpL. **E)** and **F)** Changes in the levels of the indicated mRNAs upon induction of peTrpL overproduction only. Induction time (min IPTG) is indicated.

Another direct target of rnTrpL (synonym RcsR1) is *sinI* mRNA, which encodes an autoinducer synthase. Additionally, several other mRNAs were predicted to base pair with rnTrpL, including *rplUrpmA*, which encodes ribosomal proteins L21 and L27 (Baumgardt et al., 2016). The regulation of multiple mRNAs by a single bacterial sRNA is relatively common and involves an imperfect complementarity to (most of) its targets (Storz et al., 2011; Feng et al., 2015). Frequently, these sRNAs additionally employ RNA chaperones like Hfq and ProQ to ensure efficient target binding (Santiago-Frangos and Woodson, 2018; Smirnov et al., 2016). However, the *S. meliloti* genome does not encode a ProQ homolog (Galibert et al., 2001) and rnTrpL has been reported to be an Hfq-independent sRNA (Baumgardt et al., 2016). Thus, a proteinaceous interaction partner of rnTrpL, which may enable its interaction with (specific) target mRNAs, still remained to be identified.

In addition to the well-studied ribosome-dependent attenuators in Gram-negative bacteria, recently such attenuators were also found to regulate antibiotic resistance operons of Gram-positive bacteria: upon exposure of *Bacillus* or *Listeria* to translation-inhibiting antibiotics, ribosome stalling at uORFs prevented transcription termination, thus inducing the expression of downstream resistance genes (Dar et al., 2016). The widespread occurrence and high synteny conservation of attenuator uORFs in bacteria raises the question of whether some of these leader peptides may have acquired independent functions in *trans* during evolution. This hypothesis is supported by the increasing evidence for functional small proteins encoded by sORFs shorter than 50 codons, which often are missing in current genome annotations (Storz et al., 2014). For example, in *Drosophila*, small proteins of between 11 and 32 aa in length regulate cell morphogenesis (Kondo et al., 2007) and, in *Bacillus subtilis*, the basic 29-aa protein FbpC was proposed to act as an RNA chaperone (Gaballa et al., 2008), whereas, in *E. coli,* the 31-aa protein MgtS was shown to interact with two different proteins and regulate Mg^2+^ homeostasis (Yin et al., 2018). These few examples illustrate that sORF-encoded proteins have become an important research field (Andrews and Rothnagel, 2014; Storz et al., 2014; Cabrera-Quio et al., 2016; Omasits et al., 2017; Weaver et al., 2019).

Here, we provide evidence that the *trp* attenuator-encoded leader peptide peTrpL is involved in the posttranscriptional regulation of the ribosomal genes *rplUrpmA* in *S. meliloti*. Analysis of the predicted interaction between the attenuator sRNA rnTrpL and *rplUrpmA* uncovered that peTrpL and tetracycline (Tc) are both required for an rnTrpL-mediated destabilization of *rplUrpmA*. Our results suggest that peTrpL, together with Tc or other translation-inhibiting antibiotics, redirects rnTrpL from *trpDC* to its antibiotic-dependent target *rplUrpmA*. Moreover, we found that the function of the leader peptide in *rplUrpmA* downregulation is conserved in other soil Alphaproteobacteria. In soil where antibiotic exposure is common, the downregulation of *rplUrpmA* by rnTrpL and peTrpL in a complex with an antibiotic is probably a part of an adaptation response. We propose that during evolution, the *trp* attenuator was recruited for the here described, tryptophan-independent function, because it is well suited to sense translation inhibition.

## Results

### Overproduction of the leader peptide peTrpL decreases the *rplUrpmA* mRNA level

The *rplUrpmA* operon encodes the ribosomal proteins L21 and L27 and is one of the previously predicted but not yet experimentally verified targets of the sRNA rnTrpL (Baumgardt et al., 2016). To test this prediction and potentially identify a proteinaceous partner, we constitutively overproduced the sRNA rnTrpL from the tetracycline (Tc) resistance-conferring plasmid pRK-rnTrpL (Fig. S1) in strain *S. meliloti* 2011, which has a wild type *trpLE(G)* background (Fig. 1A), and then analyzed the level of *rplUrpmA* mRNA. When compared to the empty vector control strain (EVC), the *rplUrpmA* level was increased in the constitutively overproducing strain, while, as expected, the level of the control mRNA *trpDC* was decreased and that of *trpE* was not changed (Fig. 1B).

To assess whether the increase in the *rplUrpmA* level is a primary effect, we used the gentamycin (Gm) resistance-conferring plasmid pSRKGm-rnTrpL, which allows for IPTG-inducible transcription of a rnTrpL variant named lacZ′-rnTrpL (Fig. S1). In a previous study, we could show that the sRNA lacZ′-rnTrpL is a functional riboregulator that destabilizes *trpDC* mRNA (Melior et al., 2019). However, at 10 min and 240 min (4 h) after IPTG addition to cultures of strain 2011 (pSRKGm-rnTrpL), no change in the level of *rplUrpmA* was detected, although *trpC* was downregulated, thus confirming the functionality of lacZ′-rnTrpL as a *trans*-acting sRNA (Fig. 1C).

To explore possible reasons for this unexpected result, we tested if the different vector backbone or the presence of Tc were responsible for the observed effects on the *rplUrpmA* level (Fig. 1B). We conjugated the empty vector pRK4352 into strain 2011 (pSRKGm-rnTrpL), grew the resulting two-plasmid strain 2011 (pSRKGm-rnTrpL, pRK4352) in media containing Gm and Tc, and analyzed changes in the cellular *rpmA* level at 10 min and 4 h post induction (p. i.) with IPTG. We found that, in the two-plasmid cultures, the *rpmA* mRNA level was decreased at 10 min p. i., while an increase was detected at 4 h p. i. (see Fig. 1D). This increase was similar to the increase observed in the constitutively overproducing strain 2011 (pRK-rnTrpL) (compare to Fig. 1B).

Next, we set out to overproduce only the peptide peTrpL. For this, we cloned the sORF *trpL*, with several synonymous nucleotide substitutions (to avoid RNA-based interaction with *rplUrpmA*) and without rare codons, in frame with the *lacZ* start codon in plasmid pSRKGm. We found that IPTG-induced overproduction of peTrpL (alone) was sufficient for the above described effects on *rpmA*, provided that pRK4352 was also present in the used strain and, thus, Tc was included in the growth medium (Fig. 1E and 1F). This effect of the leader peptide was specific to *rpmA*, since the levels of *trpC* and *trpE* remained unchanged (Fig. 1F).

Taken together, the results shown in Fig. 1 lead us to conclude that a decrease in the cellular level of *rpmA* mRNA (representing *rplUrpmA*) is probably a primary effect of peTrpL overproduction in strain 2011, while the increase after long-term peTrpL overproduction is a downstream effect. Furthermore, we conclude that the observed changes in the *rpmA* level may depend on Tc.

### Both peTrpL and rnTrpL are required for *rplUrpmA* downregulation

Next, we analyzed the short-term effects of rnTrpL and peTrpL on *rplUrpmA* in a *ΔtrpL* background. Strain 2011*ΔtrpL* (pSRKGm-rnTrpL, pRK4352) was grown in medium containing Tc and Gm. At 10 min p. i. of lacZ′-rnTrpL (and of peTrpL encoded by this sRNA), the *rpmA* level was decreased like in strain 2011 that still harbored the chromosomal *trpL* gene (Fig. 2A; compare to Fig. 1D). Interestingly, if 2011*ΔtrpL* (pSRKGm-peTrpL, pRK4352) was used (i.e., a *ΔtrpL* strain that only produces peTrpL upon induction), no significant change in the *rpmA* level was observed (Fig. 2A), suggesting that the negative effect on *rpmA* required the presence of both the peptide and the sRNA. To test this, we used two plasmids that (separately) conferred the sRNA function (pRK-rnTrpL-AU1,2UA; the AU1,2UA mutation in rnTrpL inactivates the sORF *trpL*, see Fig. 2B) or the peptide function (pSRKGm-peTrpL). A decrease in the *rpmA* level was detected only if both the sRNA and the peptide were produced (Fig. 2C), providing evidence that both peTrpL and rnTrpL are required to downregulate *rplUrpmA*.

**Figure 2.**
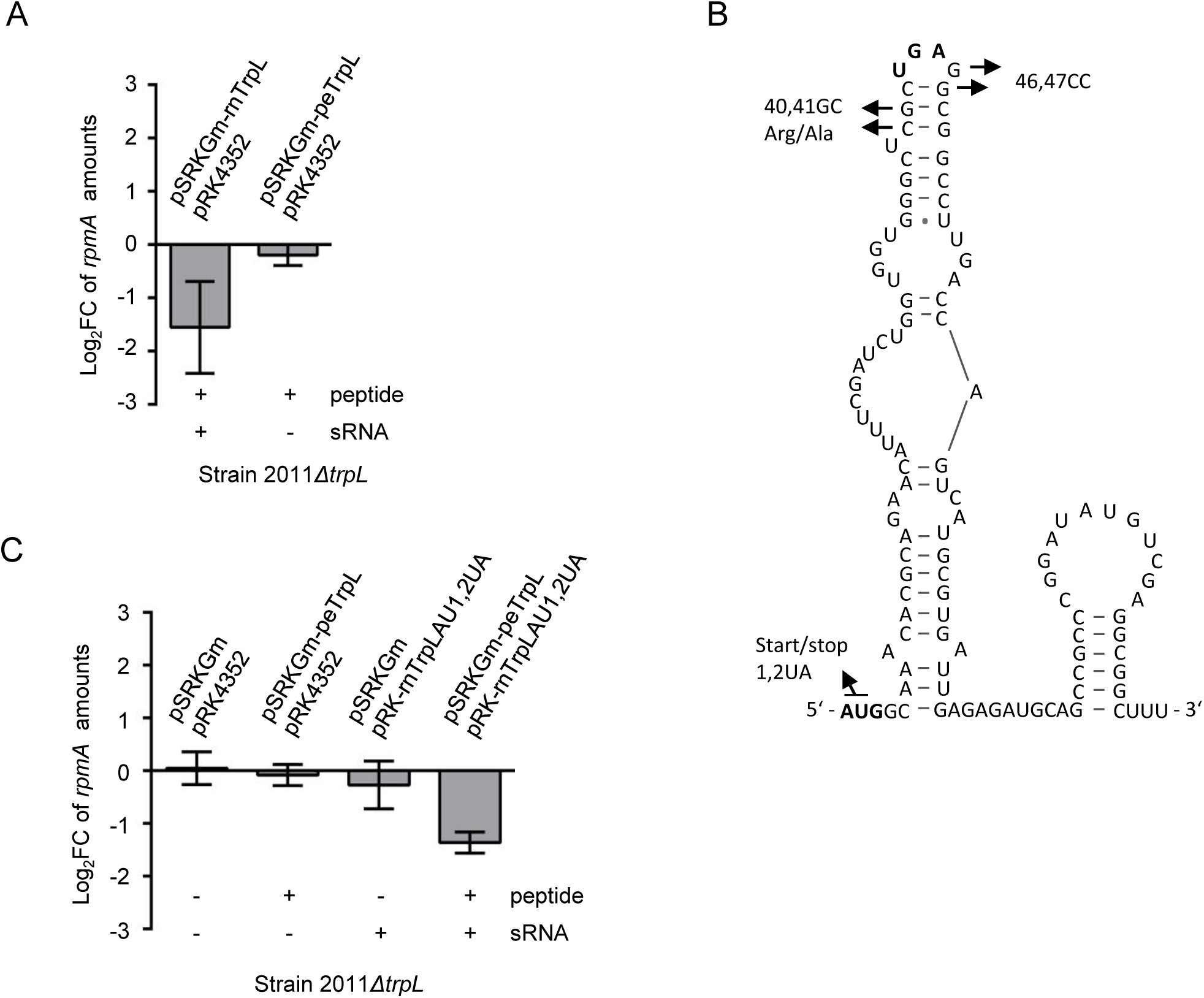
Both peTrpL and rnTrpL are required for *rplUrpmA* downregulation. **A)** and **C)** Cultures of the deletion mutant 2011*ΔtrpL* containing the indicated plasmids and grown with Gm and Tc were analyzed. Changes in the *rpmA* levels 10 min post addition of IPTG were compared to those before IPTG addition using qRT-PCR. Relevant plasmid products (sRNA and peptide) are indicated below the graphs. Each graph contains data from three independent experiments, each performed in duplicates (shown are means and standard deviations). **B)** Secondary structure of rnTrpL (according to Melior et al., 2019) with indicated mutations used in this work. The rnTrpL derivative with start codon of the *trpL* sORF exchanged for a stop codon (harbors an inactivated *trpL* sORF and exhibits the sRNA functions only) was constitutively overproduced from plasmid pRK-rnTrpL-AU1,2UA.

### Tetracycline is also required for downregulation of *rplUrpmA* by rnTrpL and peTrpL

To investigate a possible involvement of Tc in the regulation of *rplUrpmA* by rnTrpL and peTrpL, cells of strain 2011*ΔtrpL* (pSRKGm-rnTrpL, pRK4352) were washed and grown for 4 h in medium without Tc to exclude possible effects of Tc on gene expression (pRK4352 was not lost during growth without Tc; qPCR data not shown). Then, lacZ′-rnTrpL production (and, as a result, peTrpL production) were induced with IPTG (Fig. 3A). After 10 min, Tc was added and the cells were incubated for another 10 min with IPTG and Tc. RNA was isolated at 0, 10 and 20 min (Fig. 3A) and changes in the *rpmA* levels were analyzed. In parallel, control cultures were exposed to IPTG only, to Tc only or to Tc before IPTG was added. Fig. 3B shows that the *rpmA* level was only decreased if both IPTG and Tc were present in the medium. The *rpmA* level was not changed if lacZ′-rnTrpL production was induced with IPTG in Tc-free medium (t = 10 min in Fig. 3A; see the bar marked with an asterisk in Fig. 3B). Importantly, 10 min after Tc addition to this IPTG-induced culture, the *rpmA* level decreased (t = 20 min in Fig. 3A; see the bar marked with two asterisks in Fig. 3B). Since the *rpmA* decrease was similar at 10 min and 20 min after simultaneous addition of IPTG and Tc (see the two left bars in Fig. 3B), the observed change in the *rpmA* level at 10 min after Tc addition to the IPTG-induced culture (t = 20 min in Fig. 3A, the bar marked with two asterisks in Fig. 3B) can be attributed to the Tc exposure. This Tc-dependent regulation was specific to *rplUrpmA*, since the level of the *trpC* mRNA was always decreased upon lacZ′-rnTrpL induction, regardless of the presence or absence of Tc, and the level of *trpE* was essentially not changed (Fig. S2). Together, these results suggested that Tc directly contributes to the down-regulation of *rplUrpmA* by rnTrpL and peTrpL.

**Figure 3.**
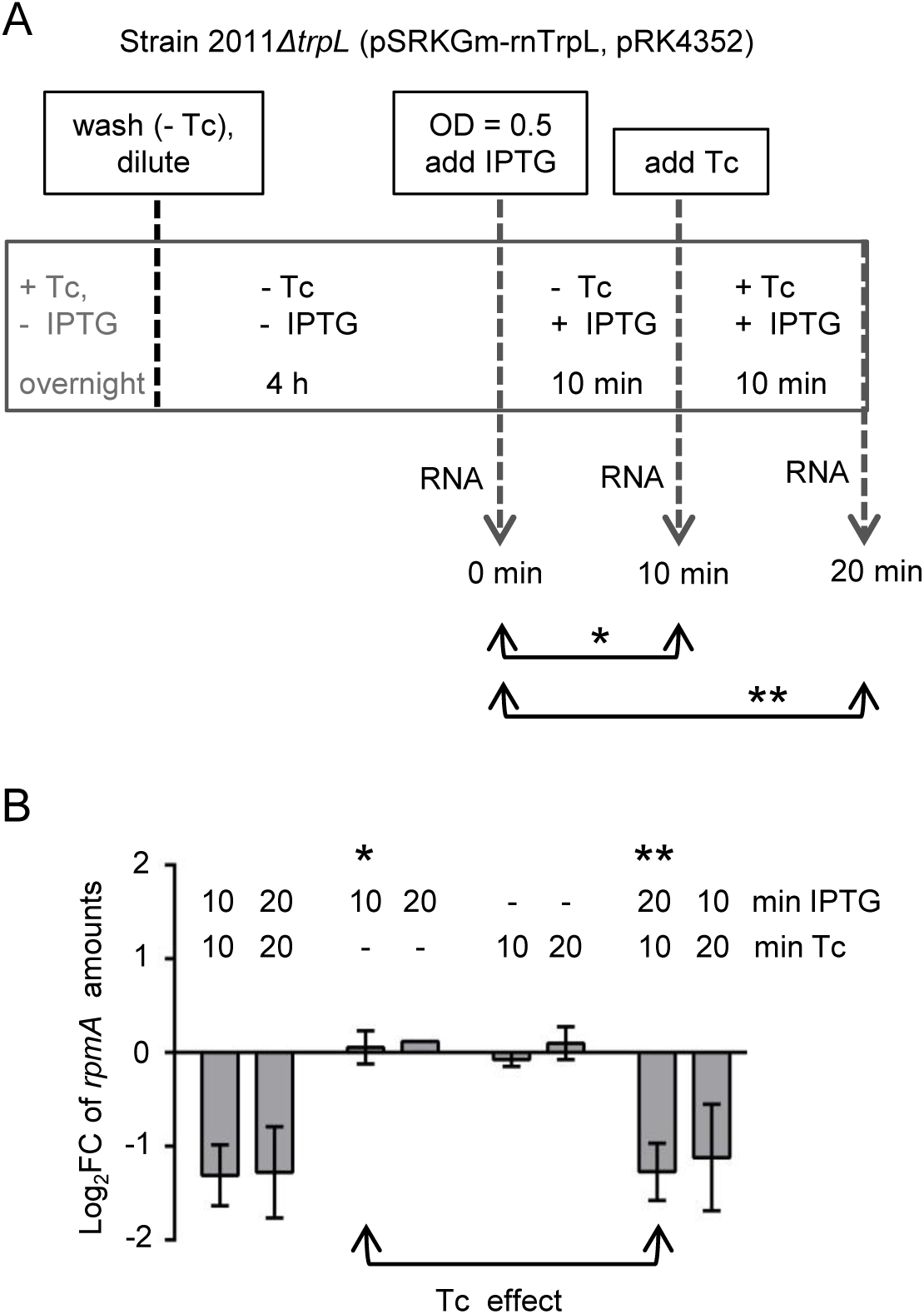
Tetracycline is also required for downregulation of *rplUrpmA* by rnTrpL and peTrpL. **A)** Schematic representation of the experiment aiming to detect a short-term effect of Tc on *rplUrpmA* expression in strain 2011*ΔtrpL* (pSRKGm-rnTrpL, pRK4352). The culture was grown first in medium with Gm and Tc. Then, the strain was grown for 4 h without Tc, IPTG was added and, 10 min later, Tc was also added. RNA was isolated at the indicated time points. The *rpmA* levels at the time points 10 and 20 min were compared to the level at the time point 0 (marked with arrows and asterisks). **B)** qRT-PCR analysis of changes in the *rpmA* level upon induction of lacZ′-rnTrpL transcription with IPTG and/or exposure to Tc. In addition to the culture treatment depicted in panel A, suitable controls were conducted: cultures were exposed to IPTG only, Tc only or to a combination of both compounds for the indicated times (min). Bars representing the comparisons of the *rpmA* levels at time point 10 min (one asterisk) and 20 min (two asterisks) to the level at time point 0 min (see panel A) are indicated. Their difference indicates a Tc effect on *rplUrpmA* expression. Shown are means and standard deviations from three independent experiments, each performed in duplicates.

### The sRNA rnTrpL base pairs with *rplU* and destabilizes *rplUrpmA* mRNA in a Tc-dependent manner

The results from the above experiments defined the conditions necessary to observe the putative primary effect of rnTrpL on *rplUrpmA*: presence of peTrpL and Tc in addition to the sRNA. This allowed us to test experimentally *in vivo* the predicted base-pairing between rnTrpL and *rplU* in *rplUrpmA* mRNA (Fig. 4A). We performed two-plasmid assays in strain 2011*ΔtrpL* using existing lacZ′-rnTrpL derivatives (transcribed from pSRKTc conferring resistance to Tc; Melior et al., 2019) and bicistronic *rplUrpmA*′*::egfp* reporter constructs (on pSRKGm conferring resistance to Gm, Fig. S1). Each *rplUrpmA*′*::egfp* construct was co-induced with wild type or mutated lacZ′-rnTrpL and fluorescence was measured at 20 min after IPTG addition. Fig. 4B shows that L27′-EGFP fluorescence derived from plasmid pSRKGm-rplUrpmA′-egfp was strongly decreased if lacZ′-rnTrpL was co-expressed, in line with the idea that the sRNA binds to *rplU* and thereby induces a reduction of *rplUrpmA*′*::egfp* mRNA levels. In contrast, sRNA derivatives harboring the GG46,47CC and CG40,41GC mutations, respectively (see Fig. 2B), did not cause this effect (Fig. 4B). Importantly, compensatory mutations in the *rplUrpmA*′*::egfp* transcript (CC/GG exchange at positions 221 and 222 of the *rplU* ORF, see Fig. 4A), which were designed to restore the base-pairing to rnTrpL-GG46,47CC, specifically restored the negative effect of the sRNA on fluorescence levels (Fig. 4B). Similarly, the introduction of a compensatory mutation, G228C, into *rplU* (Fig. 4A) could be shown to cause a down-regulation of the reporter by rnTrpL-CG40,41GC (Fig. 4B). These results validate the base-pairing between rnTrpL and *rplU* and show that even subtle changes in the base-pairing interactions may have a major impact on the downregulation of *rplUrpmA*.

**Figure 4.**
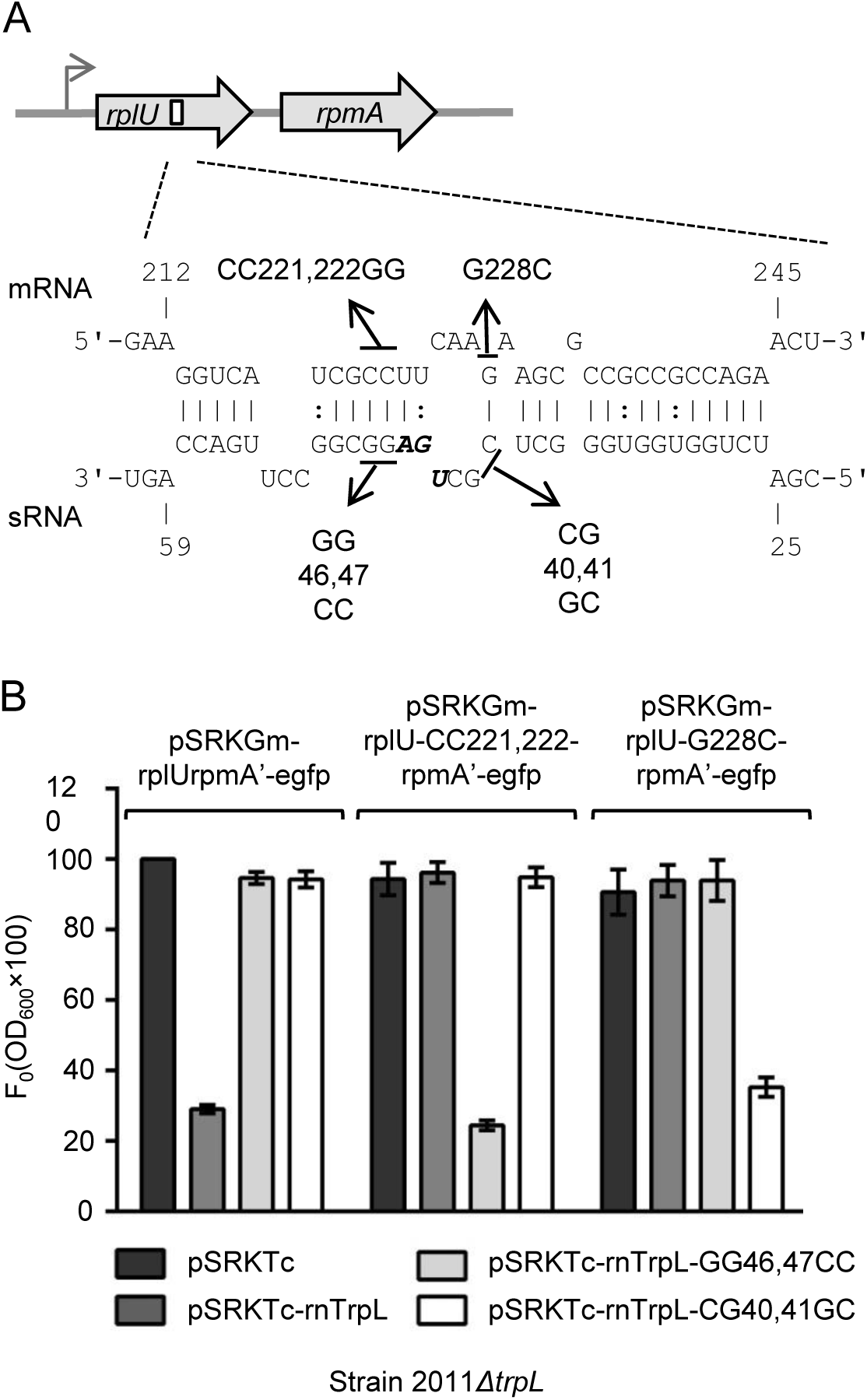
rnTrpL directly base-pairs with *rplU* to downregulate *rplUrpmA*. Scheme of the *rplUrpmA* operon and the duplex structure predicted to be formed between *rplU* (mRNA) and rnTrpL (sRNA) (ΔG = −11.54 kcal/mol). The transcription start site is indicated by a flexed arrow (Schlüter et al., 2013). The used mutations in lacZ′-rnTrpL and compensatory mutations in *rplU* are given. **B)** Analysis of possible base-pairing interactions between lacZ′-rnTrpL and the fusion mRNA *rplUrpmA*′*::egfp* in strain 2011*ΔtrpL*. The plasmids used in this experiment are indicated. Cultures were grown with Gm and Tc, and fluorescence was measured 20 min after IPTG addition. The fluorescence of strain 2011*ΔtrpL* (pSRKGm-rplUrpmA’-egfp, pSRKTc) was set to 100 % and used for normalization. Shown are means and standard deviations from three independent experiments, each performed in duplicates.

The downregulation of *rplUrpmA* by base-pairing with rnTrpL may be explained by a destabilization of the mRNA. Consistent with this hypothesis, we observed that, upon lacZ′-rnTrpL induction in strain 2011*ΔtrpL* (pSRKTc-rnTrpL) grown in medium with Tc, the half-life of *rpmA* was shortened, while that of the control mRNA *rpoB* was not changed (Fig. 5A). Since the interaction site of the sRNA rnTrpL is located in *rplU* (Fig. 4A), this result suggests that the bicistronic *rplUrpmA* mRNA is destabilized by the sRNA. To confirm this and to demonstrate that Tc is involved in the destabilization, the half-lives of *rplU* and *rpmA* were measured in strain 2011*ΔtrpL* (pSRKGm-rnTrpL) grown in medium with Gm, which was exposed to IPTG and/or Tc. The half-lives of *rplU* and *rpmA* mRNA were significantly shortened 10 min p. i. only if Tc was applied in addition to IPTG (see Fig. 5B for *rpmA* and Fig. S3 for *rplU*).

Together, the data in Fig. 4 and Fig. 5 strongly suggest that, in the presence of Tc, rnTrpL directly interacts with *rplU* and destabilizes *rplUrpmA* mRNA.

**Figure 5.**
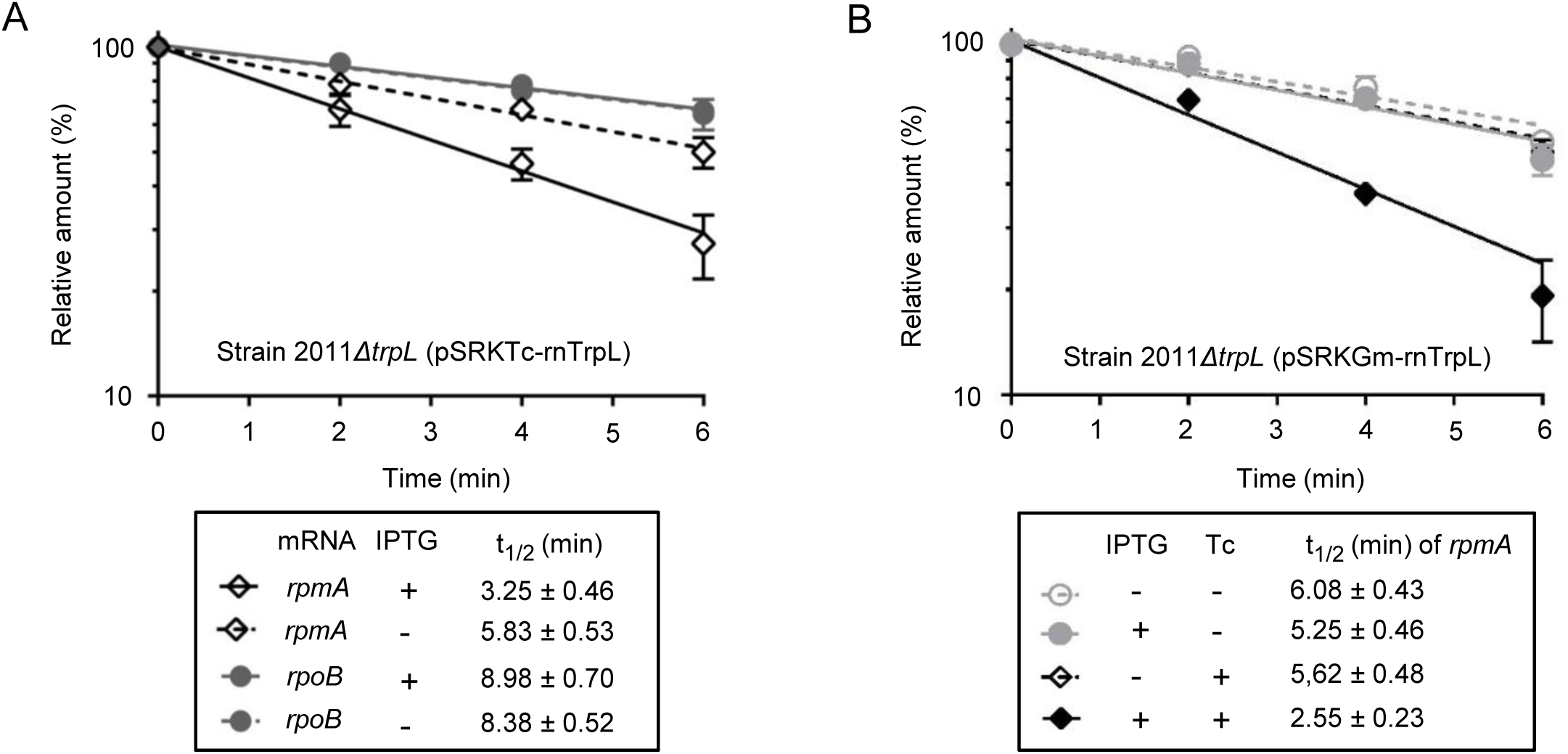
Induced rnTrpL production destabilizes *rplUrpmA* mRNA in a Tc-dependent manner. **A)** Half-lives of *rpmA* and (as a negative control) *rpoB* mRNA in strain 2011*ΔtrpL* (pSRKTc-rnTrpL) grown with Tc. 10 min after induction of lacZ′-rnTrpL production by IPTG, rifampicin (Rf) was added to stop cellular transcription. In parallel, a non-induced culture was treated with Rf. The mRNA level at time point 0 (before Rf addition) was set to 100%, the relative mRNA level values were plotted against the time, and the mRNA half-lives were calculated. Shown are the results from three independent transcription inhibition experiments. The qRT-PCRs reactions were performed in technical duplicates (means with standard deviations are indicated). **B)** Half-lives of *rpmA* in strain 2011*ΔtrpL* (pSRKGm-rnTrpL) grown with Gm and exposed to a subinhibitory Tc concentration (1.5 µg/ml) for 10 min. A culture was split in four portions, which were treated differently in respect to IPTG and Tc addition (indicated). For other descriptions see A). The *rplU* half-lives are shown in Fig. S3.

### peTrpL and rnTrpL form a tetracycline-dependent complex with *rplUrpmA*

Based on the above data, we hypothesized that peTrpL, rnTrpL, Tc and *rplUrpmA* mRNA form a complex in *S. meliloti*. To isolate this complex, we decided to produce an N-terminally triple FLAG-tagged version of TrpL (3×Flag-peTrpL) and to perform coimmunoprecipitation (CoIP) experiments. First, we tested whether 3×Flag-peTrpL is functional. We detected a decrease in the *rpmA* level at 10 min post IPTG addition to cultures of strain 2011 (pSRKGm-3×Flag-peTrpL, pRK4352) (Fig. 6A). Although this decrease was less pronounced than the decrease caused by wild type peTrpL, this result suggested that 3×Flag-peTrpL largely retained the functionality of the native peptide. However, we found that the 3×Flag-peTrpL peptide is non-functional in the deletion mutant 2011*ΔtrpL,* suggesting that the leader peptide acts as a dimer or multimer, and the tagged peptide may form functional complexes with the wild type peptide. Therefore, for the subsequent analyses, 3×Flag-peTrpL was produced in strain 2011 with the wild type *trpL* background.

**Figure 6.**
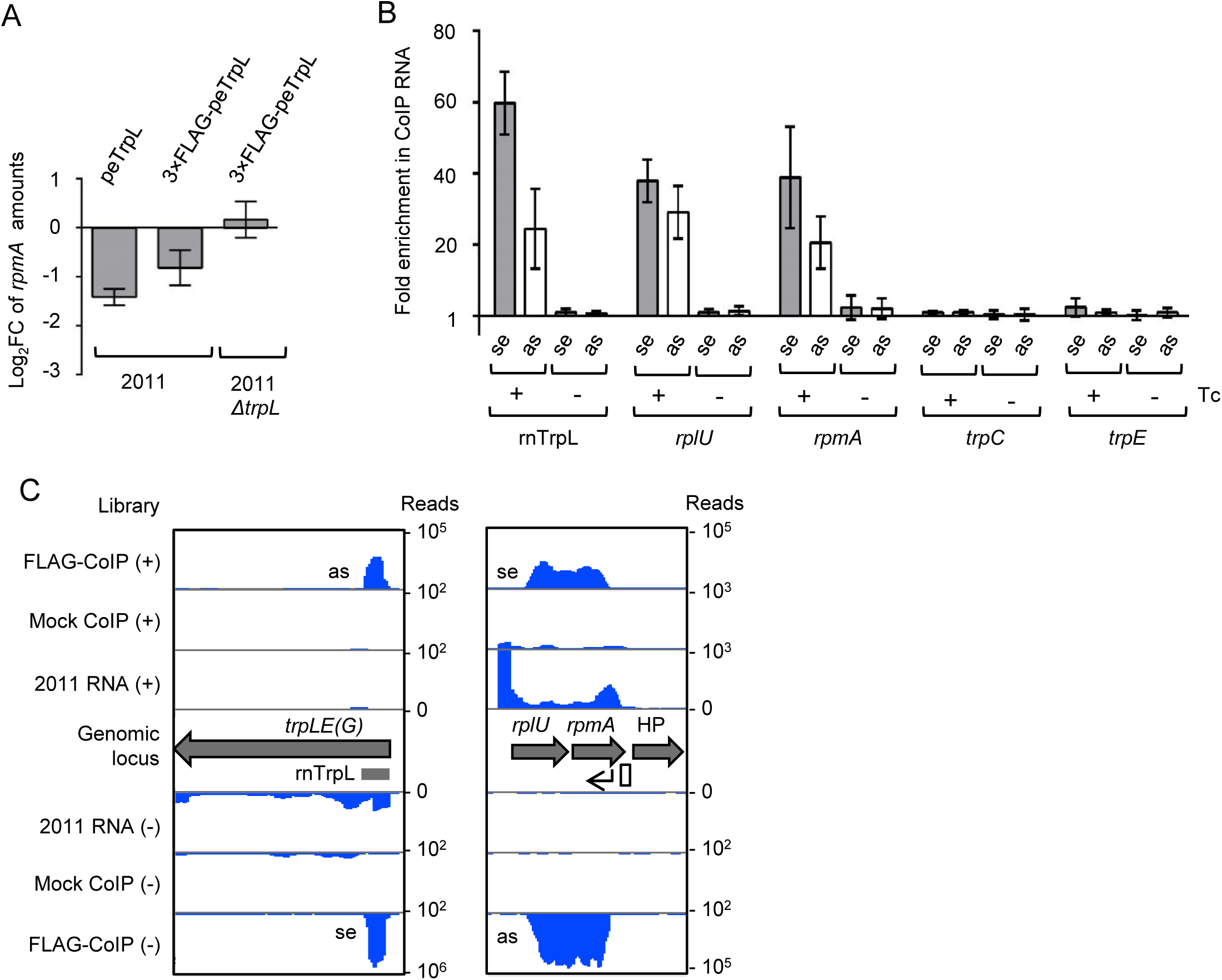
Coimmunoprecipitation with 3×FLAG-peTrpL reveals a Tc-dependent complex containing rnTrpL, *rplUrpmA* and antisense RNAs. **A)** Induced production of 3×FLAG-peTrpL down-regulates *rplUrpmA* in strain 2011 but not in the deletion mutant 2011*ΔtrpL*. Cultures of strains 2011 (pSRKGm-peTrpL, p|RK4352), 2011 (pSRKGm-3×FLAG-peTrpL, pRK4352), and 2011*ΔtrpL* (pSRKGm-3×FLAG-peTrpL, pRK4352), respectively, were grown in medium with Gm and Tc (20 µg/ml). Shown is a qRT-PCR analysis of changes in *rpmA* levels at 10 min post induction of peTrpL or 3×FLAG-peTrpL as indicated. Shown are means and standard deviations from three independent experiments, each performed in duplicates. **B)** Analysis by qRT-PCR of the indicated RNAs in the 3×FLAG-peTrpL CoIP samples from strain 2011 (pSRKGm-3×FLAG-peTrpL, pRK4352). Enrichment was calculated in comparison to mock CoIPs conducted with strain 2011 (pSRKGm-peTrpL, pRK4352). The presence of Tc in the washing buffer is indicated. se, sense RNA; as, antisense RNA. Shown are the results from three independent CoIP experiments. The qRT-PCR for each experiment was performed in duplicates (means with standard deviations are indicated). **C)** RNA-seq analysis revealed sense (se) and antisense (as) RNA of rnTrpL and *rplUrpmA* in the 3×FLAG-peTrpL CoIP samples. Only RNA retained on the beads after washing with a Tc-containing buffer was sequenced (from the FLAG-CoIP and the mock CoIP), along with total RNA of strain 2011 (2011 RNA) grown under similar conditions, but without Gm and Tc. Shown is an IGB view of mapped cDNA reads. Annotated ORFs (gray arrows) or known transcripts (gray horizontal bar) are indicated (HP, hypothetical protein). An antisense promoter downstream of *rpmA* (white rectangle) and a corresponding transcription start site (flexed arrow) are indicated.

For the CoIP, lysates were prepared at 10 min after IPTG addition to cultures of strain 2011 (pSRKGm-3×Flag-peTrpL, pRK4352) and the control strain 2011 (pSRKGm-peTrpL, pRK4352), respectively. To account for a possible role of Tc in ribonucleoprotein complex formation, the beads were divided into two fractions after incubation of the lysate with FLAG tag-specific antibodies coupled to beads. One fraction was washed with a Tc-containing buffer (2 µg/ml Tc, tenfold less than the Tc concentration in the medium), while the second fraction was washed with buffer without Tc. Then, coimmunoprecipitated RNA was purified and analyzed by qRT-PCR. The sRNA rnTrpL was strongly enriched by CoIP with the 3×FLAG-peTrpL, but only if Tc was present in the washing buffer (Fig. 6B). Similarly, *rplUrpmA* could be coprecipitated in a Tc-dependent manner, while none of the control mRNAs (*trpC, trpE*) could be coprecipitated (Fig. 6B). These data support the idea that interactions between the attenuator sRNA rnTrpL, the leader peptide peTrpL and their target mRNA *rplUrpmA* are facilitated by (or even dependent on) the presence of Tc.

We also analyzed the CoIP-RNA by RNA-seq, revealing that, in addition to rnTrpL and *rplUrpmA*, the corresponding antisense RNAs (asRNAs) were also coimmunoprecipitated (Fig. 6C). Tc-dependent enrichment of the anti-rnTrpL and anti-*rplUrpmA* transcripts by the CoIP was validated by qRT-PCR (Fig. 6B). While a transcription start site (TSS) of an anti-rnTrpL RNA was annotated previously, no TSS for the asRNA complementary to *rplUrpmA* was known (Schlüter et al., 2013). We tested the region downstream of *rplUrpmA* for antisense promoter activity using a transcriptional fusion with a reporter *egfp* mRNA (Fig. S1) and observed a strong induction of antisense transcription upon exposure to Tc. The level of the reporter *egfp* mRNA was increased 13-fold (± 4). This explains the co-purification of the anti-*rplUrpmA* RNA from a culture grown in medium with Tc, and suggests that asRNAs may play a role in a Tc-dependent regulation of *rplUrpmA* expression.

To provide additional evidence for a Tc-dependent complex between peTrpL, rnTrpL and *rplUrpmA*, we decided to produce a 5′-terminally tagged MS2-rnTrpL sRNA in *S. meliloti* and to isolate it together with its interaction partners using an MS2-MBP fusion protein coupled to amylose beads as described by Smirnov et al. (2016). First we tested the functionality of the tagged sRNA in strain 2011*ΔtrpL* (pRK-MS2-rnTrpL, pSRKGm-peTrpL). After 10 min induction of peTrpL production, a decrease in the *rpmA* level was detected (Fig. S4). In contrast, such a decrease was not detected if (instead of pRK-MS2-rnTrpL, from which MS2-rnTrpL is transcribed) the negative control plasmid pRK-MS2 encoding the MS2 aptamer only was used. Based on these data, we concluded that peTrpL is capable to downregulate *rplUrpmA* with the help of the sRNA MS2-rnTrpL (Fig. S4).

Then, lysates of 2011*ΔtrpL* (pRK-MS2-rnTrpL, pSRKGm-peTrpL) cultures were prepared at 10 min post IPTG addition and affinity chromatography was performed. The MS2-MBP amylose beads were washed in buffer containing or lacking Tc as described above, and the elution fractions were analyzed by qRT-PCR. Fig. 7A shows that, as expected, MS2-rnTrpL was strongly enriched in the elution fractions independently of the presence of Tc in the washing buffer. Compared to the amounts of rnTrpL, very low amounts of asRNA were co-purified, which probably originated from spurious transcription from the plasmid. Importantly and in line with the results shown in Fig. 6, *rplUrpmA* and its asRNA were co-purified only if washing buffer that contained Tc was used (Fig. 7A).

**Figure 7.**
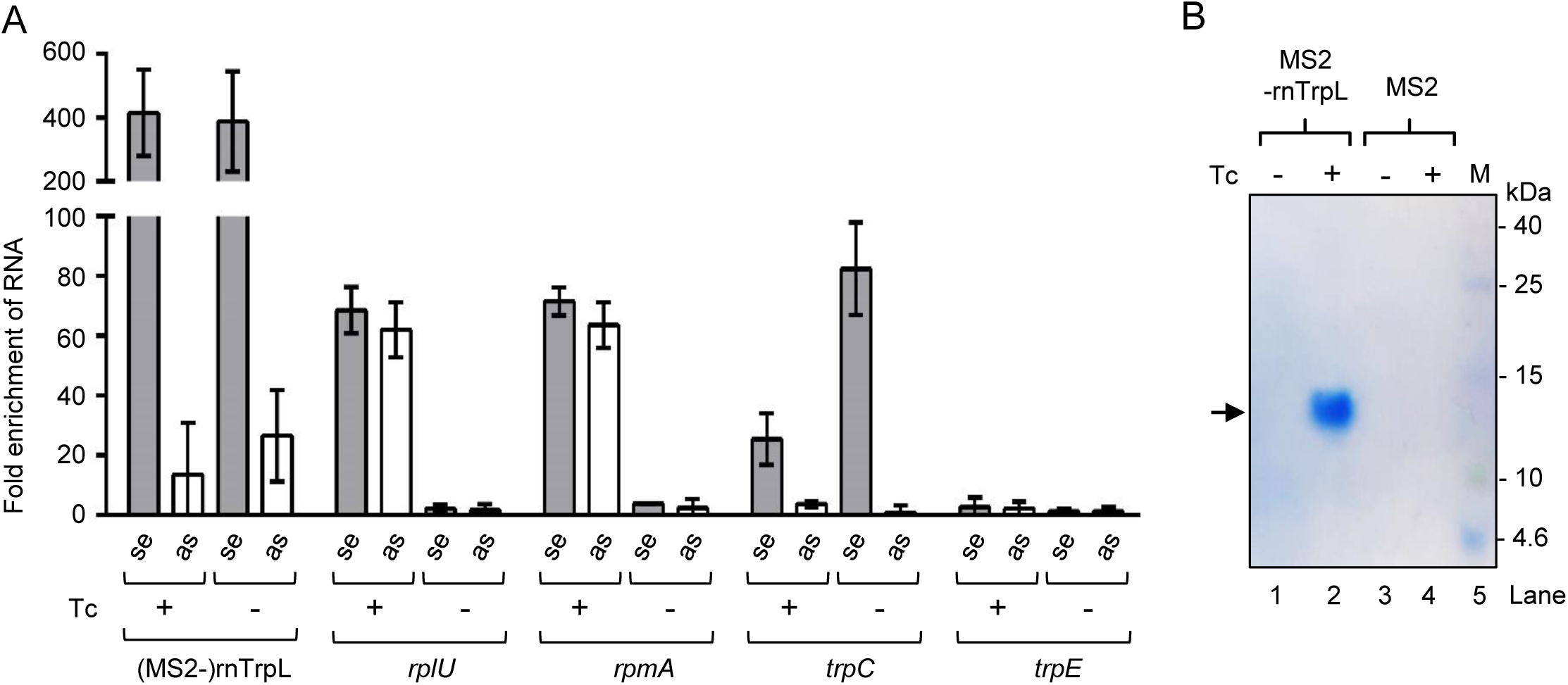
Affinity purification of MS2-rnTrpL confirms a Tc-dependent ribonucleoprotein complex (ARNP). **A)** Analysis by qRT-PCR of the indicated RNAs in the elution samples of MS2-MBP affinity chromatography of strain 2011 (pSRKGm-3×FLAG-peTrpL, pRK-MS2-rnTrpL) producing the aptamer-tagged sRNA MS2-rnTrpL. Enrichment was calculated in comparison to the elution samples of similar chromatography of strain 2011 (pSRKGm-3×FLAG-peTrpL, pRK-MS2) producing only the MS2 aptamer. For other descriptions see Fig. 6B. **B)** Tricine-SDS-PAGE analysis of the MS2-rnTrpL and MS2 samples described in A). Migration behavior of protein standards is indicated (in kDa). The 3×FLAG-peTrpL band is marked with an arrow.

As controls, *trpC* and *trpE* were analyzed. Concerning the Tc-independent target *trpC* (representative of *trpDC*), we made the following observations: i) relatively low amounts of *trpC* were co-purified when Tc was present in the washing buffer; ii) the amount of co-purified *trpC* was significantly higher when buffer without Tc was used in the washing procedure; iii) no asRNA complementary to *trpC* was co-purified (Fig. 7A). The last observation suggests that asRNA is not involved in the *trpDC* regulation by rnTrpL. Further, the increased co-purification of *trpC* upon washing without Tc suggests that, during the washing step (performed in batch), *rplUrpmA* and peTrpL were released from MS2-rnTrpL and, concomitantly, the remaining *trpD* present in the residual diluted lysate was bound. This suggestion was corroborated *in vitro* (see below and Fig. 8D). Finally, *trpE* mRNA (a negative control that does not interact with rnTrpL) was not co-purified, showing the specificity of target mRNA copurification with MS2-rnTrpL (Fig. 7A).

**Figure 8.**
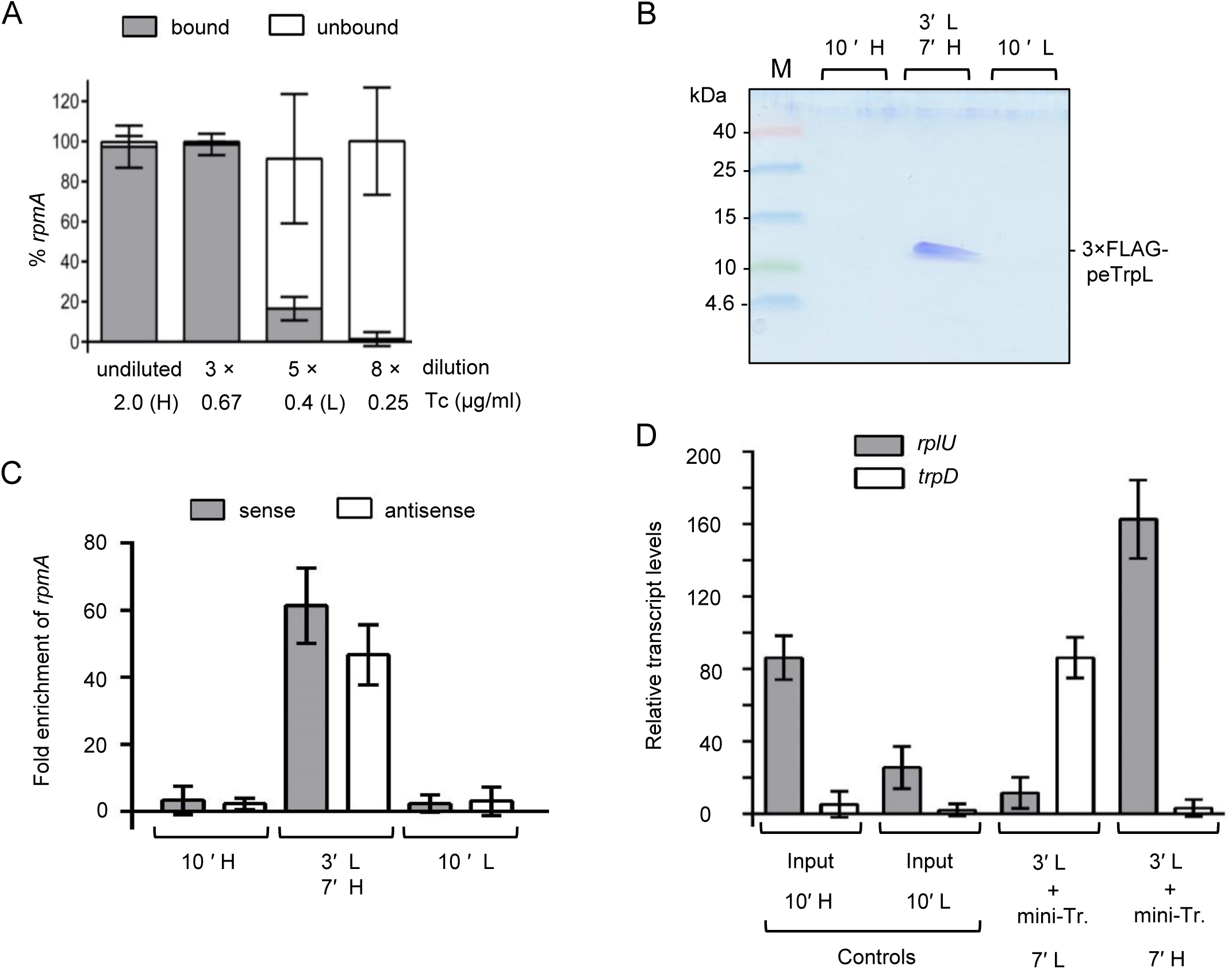
Increased Tc concentration cause ARNP assembly and redirection of rnTrpL from *trpD* to *rplU.* ARNP complexes containing only the wild type peTrpL were purified from *S. meliloti* containing pRK-MS2-rnTrpL, eluted in buffer with 2 µg/ml Tc (H, High-Tc), and subjected to dissociation and reassociation. **A)** The eluted ARNP complex was serially diluted with a buffer without Tc as indicated, and then pull-down with MS2-MBP-beads was performed. The supernatant (unbound *rpmA*) and pellet (*rpmA* bound to MS2-rnTrpL) were compared by qRT-PCR. The complex largely dissociates at 0.4 µg/ml Tc (L, Low-Tc). For **B)** and **C),** the eluted ARNP complex was amended with synthetic 3×FLAG-peTrpL and divided in three portions that were treated differently: 1) 10′ H, the sample was incubated for 10 min at High-Tc. 2) 3′ L, 7′ H, the sample was diluted 5× in buffer without Tc and incubated for 3 min at Low-Tc; then the Tc concentration was raised and the sample was incubated for 7 min at High-Tc. 3) 10′ L, the sample was diluted 5× in buffer without Tc and incubated for 3 min at Low-Tc; then corresponding volume of ethanol (Tc solvent) was added, without raising the Tc concentration, and the sample was incubated for additional 7 min at Low-Tc. **B)** The treated samples were subjected to MS2-MBP affinity chromatography and the elution fractions were analyzed by Tricine-SDS-PAGE. A representative Coomassie stained gel is shown. **C)** The treated samples were subjected to CoIP with FLAG-specific antibodies and coimmunoprecipitated RNA was analyzed by qRT-PCR with *rpmA* specific primers. **D)** To an ARNP sample that was diluted 5× in buffer without Tc, incubated for 3 min, and mini-*rplU* and mini-*trpC* transcripts were added. The sample was split and the one half was kept at Low-Tc, while to the second half Tc was added to achieve High-Tc conditions. After seven minutes, MS2-MBP affinity chromatography was performed and the transcripts co-purified with MS2-rnTrpL were analyzed by qRT-PCR. For details see B) and C). All graphs show are means and standard deviations from three independent experiments, each performed in duplicates.

To test for the presence of the leader peptide in the Tc-dependent MS2-rnTrpL-complex, we took advantage of the 3×FLAG-peTrpL (4.5 kDa) because we were not able to detect the very small peTrpL (1.8 kDa) by SDS-PAGE. Strain 2011 (pRK-MS2-rnTrpL, pSRKGm-3×FLAG-peTrpL) was used for MS2-MBP affinity chromatography, and the elution fractions were analyzed by a Tricine-SDS-PAGE. We detected a small protein migrating at approximately 13 kDa, which was copurified with MS2-rnTrpL only if the Tc-containing buffer was used (Fig. 7B, lane 2). This band was identified by Western blot analysis (Fig. S5) and mass spectrometry (Table S1) as aberrantly migrating 3×FLAG-peTrpL. The lack of 3×FLAG-peTrpL in the MS2-rnTrpL sample that was washed with buffer without Tc (Fig. 7B, lane 1) confirmed that Tc is required for the copurification of the leader peptide with the attenuator sRNA. Other proteins were not detected in the purified complex (Fig. S6).

Taken together, the data shown in Fig. 6 and Fig. 7 strongly suggest that peTrpL, rnTrpL and *rplUrpmA* form an <u>a</u>ntibiotic-dependent ribo<u>n</u>ucleo<u>p</u>rotein complex (ARNP) that also contains asRNA.

### Tetracycline causes ARNP assembly and redirection of rnTrpL from *trpD* to *rplU*

To demonstrate that Tc mediates the ribonucleoprotein complex formation, we studied *in vitro* complex disassembly by diluting an ARNP sample (and thus lowering the concentration of Tc, among others), and reassembly by raising the Tc concentration in the diluted sample. First the disassembly of ARNP purified via MS2-rnTrpL from strain 2011*ΔtrpL* (pRK-MS2-rnTrpL, pSRKGm-peTrpL) was analyzed. Five-fold dilution of the sample resulted in a profoundly reduced interaction between MS2-rnTrpL and *rplUrpmA* (Fig. 8A). Therefore, this dilution, which decreased the Tc concentration from 2 µg/ml (High-Tc, H) to 0.4 µg/ml (Low-Tc, L), was used for complex disassembly in subsequent experiments.

The reassembly was tested by adding 5 µg synthetic 3×FLAG-peTrpL to an ARNP sample that contained 2.4 µg protein. The sample (in High-Tc buffer) was divided in three portions. Two control portions, which were diluted 5-fold, were kept at High-Tc or Low-Tc conditions for 10 min (10′ H and 10′ L, respectively). The third portion was first subjected to complex disassembly (5-fold dilution and incubation at Low-Tc for 3 min; 3′ L) and then the Tc concentration was raised to reassemble ARNPs (incubation at High-Tc for 7 min; 7′ H). The three samples were analyzed by MS2-MBP affinity chromatography (Fig. 8B) and by CoIP with FLAG-specific antibodies (Fig. 8C). When MS2-MBP affinity chromatography was conducted, the 3×FLAG-peTrpL was co-purified with MS2-rnTrpL only from the sample that was subjected to disassembly and reassembly, but not in the control samples (Fig. 8B). Consistently, Fig. 8C shows that *rpmA* mRNA was also coimmunoprecipitated with 3×FLAG-peTrpL only from the sample in which complex disassembly and reassembly took place. This result shows that, during reassembly, the synthetic peptide entered the complex. Additional reassembly experiments are shown in Fig. S7. Together, Fig. 8A, 8B and 8C confirm that Tc is crucial to the ARNP formation.

Next, we tested *in vitro* the impact of Tc on the substrate preference of rnTrpL. For this we designed mini-*rplU* and mini-*trpD in vitro* transcripts encompassing the respective rnTrpL-binding sites (for predicted secondary structures see Fig. S8). An ARNP complex purified from strain 2011 (pRK-MS2-rnTrpL) was used. It contained *rplU* (representing *rplUrpmA*) which was released under Low-Tc conditions (see the controls in Fig. 8D). The ARNP complex was disassembled at Low-Tc for 3 min, and then the two mini-transcripts were added in a ratio of 1:1 (300 ng of each transcript). The sample was split: one half was kept at Low-Tc, while the Tc concentration of the second half was raised to reconstitute ARNPs under High-Tc conditions. After seven minutes, MS2-MBP affinity chromatography was performed and the transcripts co-purified with MS2-rnTrpL were analyzed by qRT-PCR. Fig. 8D shows that under Low-Tc conditions mostly mini-*trpD* was co-purified (see bars [3′ L, + mini-Tr., 7′ L] in Fig. 8D). In contrast, when the Tc concentration was raised, exclusively *rplU* (mini-*rplU* and probably endogenous *rplU*) was co-purified (see bars [3′ L, + mini-Tr., 7′ H] in Fig. 8D). This result supports the view that Tc (with the help of peTrpL) redirects the sRNA rnTrpL from targets such as *trpDC* to *rplUrpmA*.

### ARNP reconstitution reveals a four-component core complex

To analyze the ARNP formation in more detail, we conducted reconstitution experiments with synthetic components only. A 1:1 mixture of synthetic peTrpL and 3×FLAG-peTrpL peptides was incubated with *in vitro* transcripts of MS2-rnTrpL and/or mini-*rplU*. The samples were supplemented with 2 µg/ml Tc or were incubated without Tc, and CoIP with FLAG-specific antibodies was performed. The coimmunoprecipitated RNAs (RNAs interacting with 3×FLAG-peTrpL under the particular conditions) were analyzed by qRT-PCR. Fig. 9A shows that only when all four components, 1) the sRNA MS2-rnTrpL, 2) the target RNA mini-*rplU*, 3) peptide (peTrpL and 3×FLAG-peTrpL), and 4) Tc were present in the mixture, both transcripts were coimmunoprecipitated with the peptide. The lack of one of the components (Tc, MS2-rnTrpL or mini-*rplU*) abolished complex formation with the peptide. The above result shows that the four components sRNA, target RNA, peptide and Tc are necessary and sufficient for the formation of a core ARNP.

**Figure 9.**
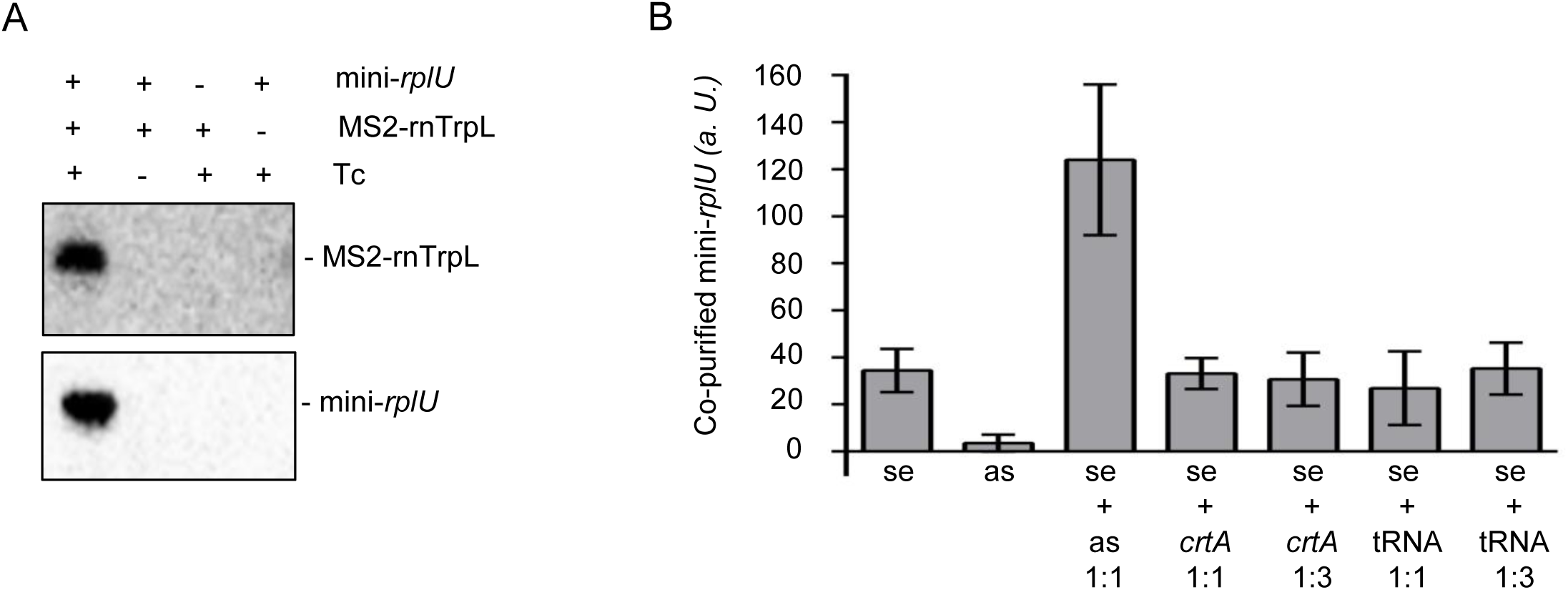
ARNP reconstitution: core complex and accessory asRNA. **A)** 50 ng synthetic peTrpL, 50 ng synthetic 3×FLAG-peTrpL, 100 ng MS2-rnTrpL *in vitro* transcript, 100 ng mini-*rplU* transcript and 2 µg/ml Tc were used. All 250 µl reaction mixtures contained the peptides. The presence of each of the other components is indicated. After 20 min incubation at 20 °C, CoIP with FLAG-directed antibodies was performed. Presence of Tc in the washing buffer corresponded to the incubation conditions. Coimmunoprecipitated RNA was analyzed by Northern blot hybridization. Detected transcripts are indicated. Results from a representative experiment are shown. **B)** 50 ng synthetic peTrpL, 50 ng synthetic 3×FLAG-peTrpL and 100 ng MS2-rnTrpL *in vitro* transcript per reconstitution reaction (final volume of 50 µl) were mixed in a buffer containing 2 µg/ml Tc. Seven reconstitution reactions were performed in parallel. Addition of 100 ng mini-*rplU* transcript (se), 100 ng corresponding antisense transcript (as) or both se and as (each 100 ng) in the first three samples is indicated. In the other four samples, unrelated *crtA in vitro* transcript (100 ng or 300 ng) or yeast tRNA (100 ng or 300 ng) was added together with the mini-*rplU* (100 ng). After 20 min incubation at 20 °C, MS2-MBP affinity chromatography was performed. Tc was included in the washing buffer. Relative levels of mini-*rplU* (se) were determined by qRT-PCR. MS2-rnTrpL was used for normalization. Shown are means and standard deviations from three independent reconstitution experiments. Each qRT-PCR was performed in technical duplicates.

We also tested whether an anti-*rplU* transcript would have an influence on the ARNP formation. Fig. 9B shows that addition of the asRNA to the reconstitution mixture strongly increased the amount of mini-*rplU* that was co-purified with MS2-rnTrpL, while unrelated, control RNAs had no effect. Thus, the asRNA increases the efficiency of ARNP formation *in vitro*. This result suggests that, in *S. meliloti*, the Tc-induced anti-*rplUrpmA* may have a supportive role in *rplUrpmA* downregulation upon exposure to Tc.

### The *trp* attenuator responds to Tc exposure by generation of the sRNA rnTrpL

The above results show that the products of the *trp* attenuator, the sRNA rnTrpL and the leader peptide peTrpL, are involved in the posttranscriptional downregulation of *rplUrpmA* upon exposure to Tc. What could be the functional relevance of such a regulatory pathway? Given that, in this pathway, ribosomal genes are regulated and Tc inhibits translation (Nelson and Levy, 2011; Chopra and Roberts, 2001), it is tempting to speculate that it represents a response to translation inhibition. We propose that the *trp* attenuator got involved in this regulation because it is well suited to sense stalled ribosomes.

As mentioned in the introduction, it is known that, in *E. coli*, pausing of the ribosome in the first half of the uORF *trpL* results in transcription termination (Zurawski et al., 1978) and thus in the release of the attenuator sRNA. Based on this, we hypothesized that, in addition to sensing the availability of charged tRNA^Trp^, the *trp* attenuator senses translation inhibition as an independent stimulus and responds by the generation of rnTrpL. Thus, even under conditions of Trp limitation, when *trpLE(G)* should be co-transcribed, exposure to Tc would arrest ribosomes at the start of *trpL* in new transcripts, and their transcription would be terminated. The released rnTrpL could then interact with Tc and peTrpL (which should be present in the cell because it is translated at least once by each transcription event) and downregulate *rplUrpmA.* Thus, we propose that, upon exposure to Tc, the *trp* attenuator is used by the cell to supply the sRNA rnTrpL independently of the Trp availability.

To test this hypothesis, we used strain 2011*ΔtrpC,* in which the attenuation can be easily assessed (Bae and Crawford, 1990; Melior et al., 2019). After 4 h of growth under Trp-limiting conditions (minimal medium supplemented with 2 µg/ml Trp), rnTrpL was essentially non-detectable in the Northern hybridization (Fig. 10A, lanes 2 and 3), a result that is consistent with the expected *trpLE(G)* cotranscription. Addition of 1.5 µg/ml Tc to this culture (the minimal inhibitory Tc concentration under these conditions was 2 µg/ml) resulted in increased rnTrpL levels (Fig. 10A, lanes 4 and 5). This is in line with the proposed transcription termination as a consequence of ribosome pausing at the first *trpL* codons. Accordingly, removal of Tc from the medium led again to a decrease in the rnTrpL levels (compare lanes 7 and 8 in Fig. 10A). These results support the idea that the *trp* attenuator can sense translation inhibition as an independent signal. They prove that, upon translation inhibition, the regulatory sRNA rnTrpL is generated even under Trp limiting conditions, in line with its Trp-independent role in *rplUrpmA* destabilization (see Fig. 10C).

**Figure 10.**
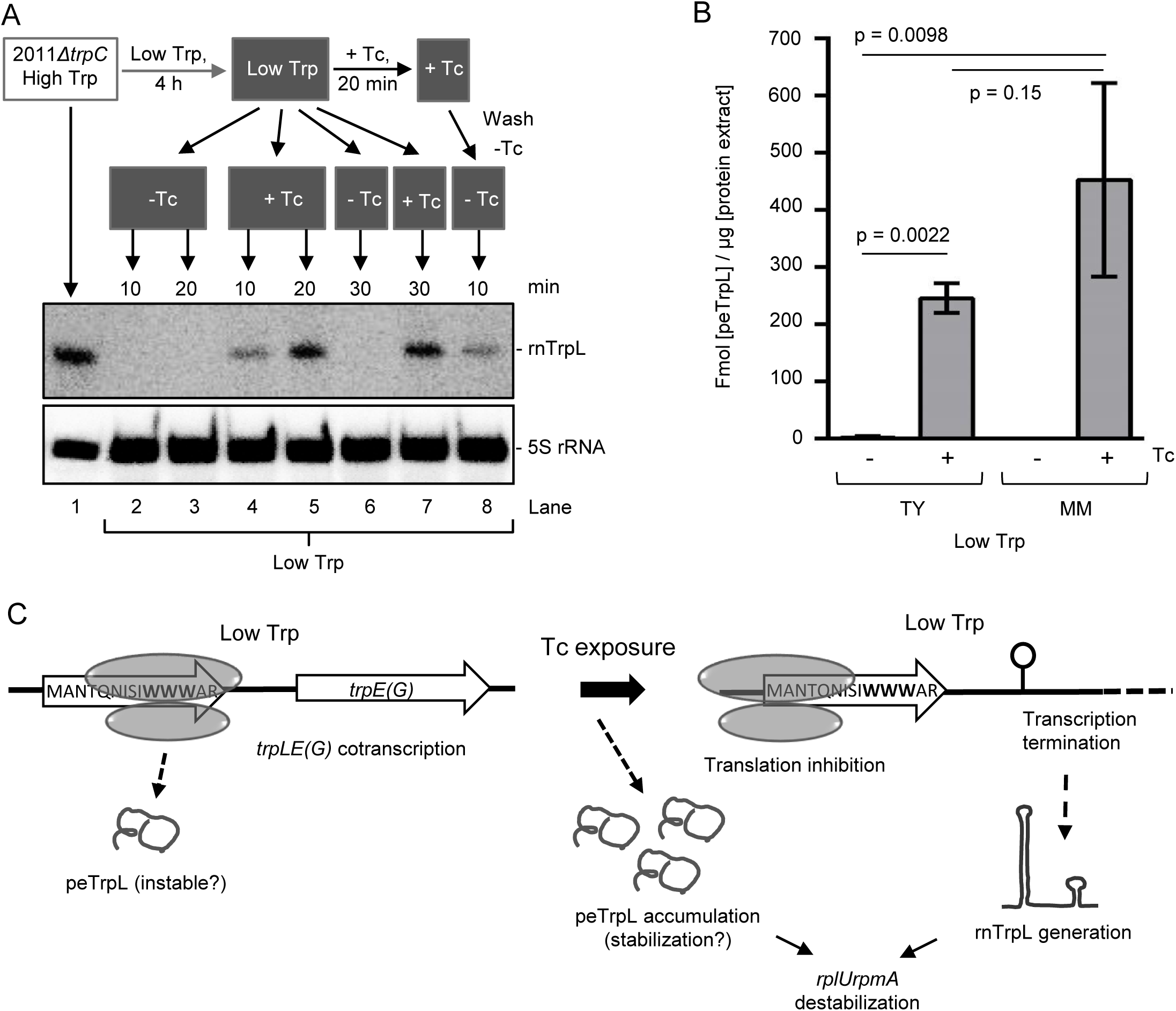
Generation of rnTrpL and accumulation of peTrpL upon exposure of *S. meliloti* to sub-inhibitory concentration of Tc. **A)** Northern blot analysis of strain 2011*ΔtrpC* shows the generation of rnTrpL under conditions of Trp insufficiency, upon exposure to Tc. Schematic representation of the experiment is shown above the hybridization panels. High Trp conditions, 20 µg/ml Trp in the minimal medium (MM); Low Trp conditions, 2 µg/ml Trp in the MM. + Tc, Tc was added to a final concentration of 1.5 µg/ml. - Tc, corresponding volume of the solvent ethanol was added. 30 µg total RNA was loaded in each lane, except for lane 1 in which 10 µg RNA was loaded. First, a probe directed against rnTrpL was used and, then, the membrane was re-hybridized with the 5S rRNA-specific probe (loading control). Detected RNAs are indicated. **B)** Mass spectrometry analysis reveals strong accumulation of peTrpL in strain 2011 upon 10 min exposure to Tc (1.5 µg/ml). Lysates of cultures grown in rich TY medium or in MM supplemented with 2 µg/ml Trp were analyzed. As indicated, the cultures were exposed to Tc or were not exposed (the solvent ethanol only was added). Shown are means and standard deviations from three independent experiments. p-values for students t-tests are given. **C)** Graphic summary of the results shown in A) and B). Under conditions of Trp insufficiency (Low Trp), peTrpL is slowly translated and probably rapidly degraded. Upon exposure to 1.5 µg/ml Tc, peTrpL accumulates, probably due to its stabilization. The (partial) translation inhibition by leads to transcription termination between *trpL* and *trpE(G)* and generation of rnTrpL. Thus, even under conditions of Trp shortage, *rplUrpmA* can be downregulated by rnTrpL and peTrpL in response to Tc exposure.

### The leader peptide peTrpL strongly accumulates upon exposure to Tc

According to our data, downregulation of *rplUrpmA* in the presence of Tc is mediated by rnTrpL and peTrpL (in combination). However, nothing is known about the abundance of peTrpL under different conditions, especially upon exposure of bacteria to Tc. Therefore, we used mass spectrometry to determine the amount of peTrpL in bacterial lysates. As a standard, we used a heavy synthetic peptide (with a heavy C-terminal arginine). *S. meliloti* 2011 was grown in rich TY medium or in minimal medium with 2 µg/ml Trp. At OD_600_ = 0.5, the cultures were divided into two halves. To one of them, 1.5 µg/ml Tc was added, and the second (control) sample was supplemented with the corresponding amount of solvent (ethanol). Cells were harvested 10 min after this treatment and the endogenous peTrpL peptide was quantified (Fig. 10B). In the control TY cultures, the peTrpL amount was very low (2,3 fmol/µl lysate), while in the Tc-treated cultures the amount was 100-fold higher. In the control cultures grown in minimal medium, the peptide was under the limit of detection, but the exposure to Tc resulted in its strong accumulation, in line with its function under these conditions (Fig. 10B and Fig. 10C).

### Several translation inhibitors support the complex of peTrpL, rnTrpL and *rplUrpmA*

The data presented above strongly suggest that *rplUrpmA* downregulation by rnTrpL and peTrpL is a response to translation inhibition by Tc. They prompted us to test whether downregulation is also triggered by other translation inhibiting antibiotics such as erythromycin (Em), chloramphenicol (Cl) and kanamycin (Km). As a negative control, the transcription inhibitor rifampicin (Rf) was included, and Tc was used as a positive control. The antibiotics were added in subinhibitory concentrations to cultures of the parental strain 2011 and the deletion mutant 2011*ΔtrpL* and, 10 min post addition, changes in the levels of *rpmA* and the control mRNA *trpE(G)* were analyzed by qRT-PCR. Fig. 11A shows that, in strain 2011, the *rpmA* level was decreased when Tc, Em, Cl or Km were applied, but not upon addition of Rf. In contrast, an *rpm*A decrease was not detected in strain 2011*ΔtrpL* after exposure to the translation inhibitors. Further, the mRNA *trpE(G)* was not significantly affected in both strains (Fig. 11A). These results support the critical role of the *trp* attenuator (and its *trans*-acting products) in *rplUrpmA* downregulation upon exposure to translation inhibitors.

**Figure 11.**
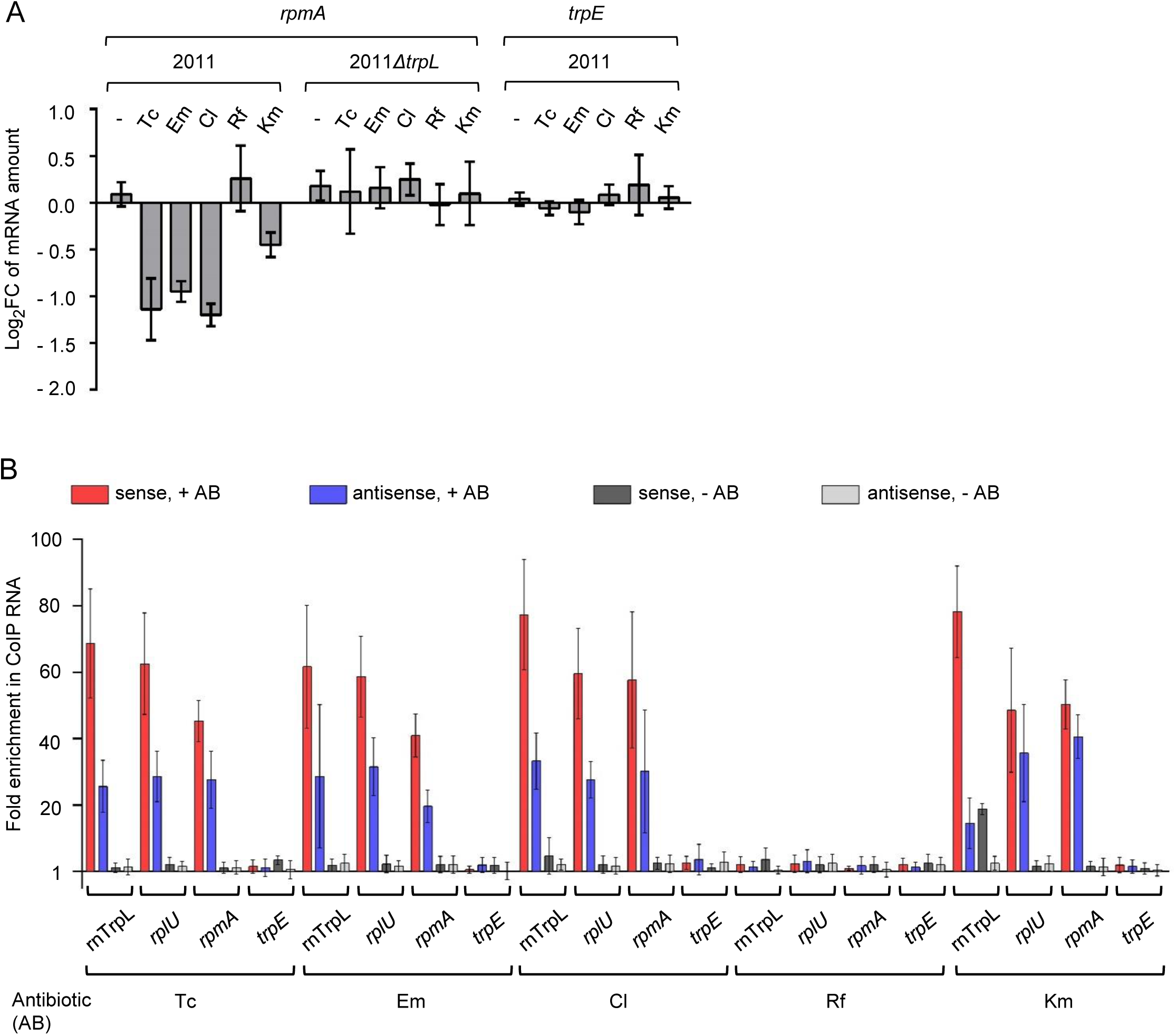
Several translation-inhibiting antibiotics down-regulate *rplUrpmA* and promote ARNP formation. **A)** qRT-PCR analysis of changes in the *rpmA* level 10 min after addition of the indicated antibiotics at subinhibitory concentrations to cultures of strain 2011 or 2011*ΔtrpL*: 1.5 µg/ml Tc, 27 µg/ml Em, 9 µg/ml Cl, 3 µg/ml Rf, 45 µg/ml Km. *trpE*, negative control. **B)** Enrichment analysis of the indicated RNAs in the 3×FLAG-peTrpL CoIP samples from strain 2011 (pSRKGm-3×FLAG-peTrpL), in comparison to the control, mock CoIP conducted with strain 2011 (pSRKGm-peTrpL). Bacterial cultures were grown in medium with Gm. CoIP was performed 10 min after addition of IPTG and one of the indicated antibiotics (the concentrations are specified in A). The same antibiotic concentration was present in the washing buffer (+ AB), or buffer without antibiotic was used (- AB). Shown are the results from three independent CoIP experiments. The qRT-PCR for each experiment was performed in duplicates (means with standard deviations are indicated).

Next, we tested whether (similar to Tc) the translation inhibiting antibiotics Em, Cl and Km support the formation of complexes involving peTrpL, rnTrpL and *rplUrpmA*. For this, strain 2011 (pSRKGm-3×FLAG-peTrpL) was grown in Gm-containing medium. Simultaneously to the induction of 3×FLAG-peTrpL production by adding IPTG, one of the antibiotics Tc, Em, Cl, Km or Rf was added. 10 min later, a CoIP with 3×FLAG-peTrpL was conducted, and one half of the beads of each CoIP was washed with a buffer containing the respective antibiotic, while the other half was washed with a buffer lacking the antibiotic. qRT-PCR analysis revealed that rnTrpL and *rplUrpmA* but not *trpE(G)* were coimmunoprecipitated in the presence of Em, Cl, Km and the positive control Tc, but not in the presence of Rf (Fig. 11B). Similar to the data shown before for cultures containing a Tc-resistance plasmid and grown at selective Tc concentration (20 µg/ml) (Fig. 6B), the experiments using subinhibitory concentrations of the various translation-inhibiting antibiotics revealed that an asRNA complementary to *rplUrpmA* was co-purified (Fig. 11B).

Together, these results suggest that (similar to Tc) the translation inhibitors Em, Cl and Km promote the formation of an ARNP that is involved in *rplUrpmA* destabilization. Thus, the posttranscriptional downregulation of *rplUrpmA* by peTrpL and rnTrpL is probably a general response to translation-inhibiting antibiotics.

### Conservation of rnTrpL and peTrpL functions in other bacteria despite sequence heterogeneity

To determine functionally important residues in peTrpL of *S. meliloti*, we performed alanine scanning mutagenesis and tested the functionality of the mutagenized peptides in strain 2011. Compared to the wild type peptide peTrpL, Ala substitutions of Thr4, Ser8 and Trp12, respectively, had the strongest impact. Induced production of these mutated peptides for 10 min led to increased (rather than decreased) *rpmA* levels (Fig. 12A). The Ala substitutions of the Trp10 and Trp11 residues abolished the effect of peTrpL overproduction, suggesting that all three Trp residues are important for the function of peTrpL (Fig. 12A).

**Figure 12.**
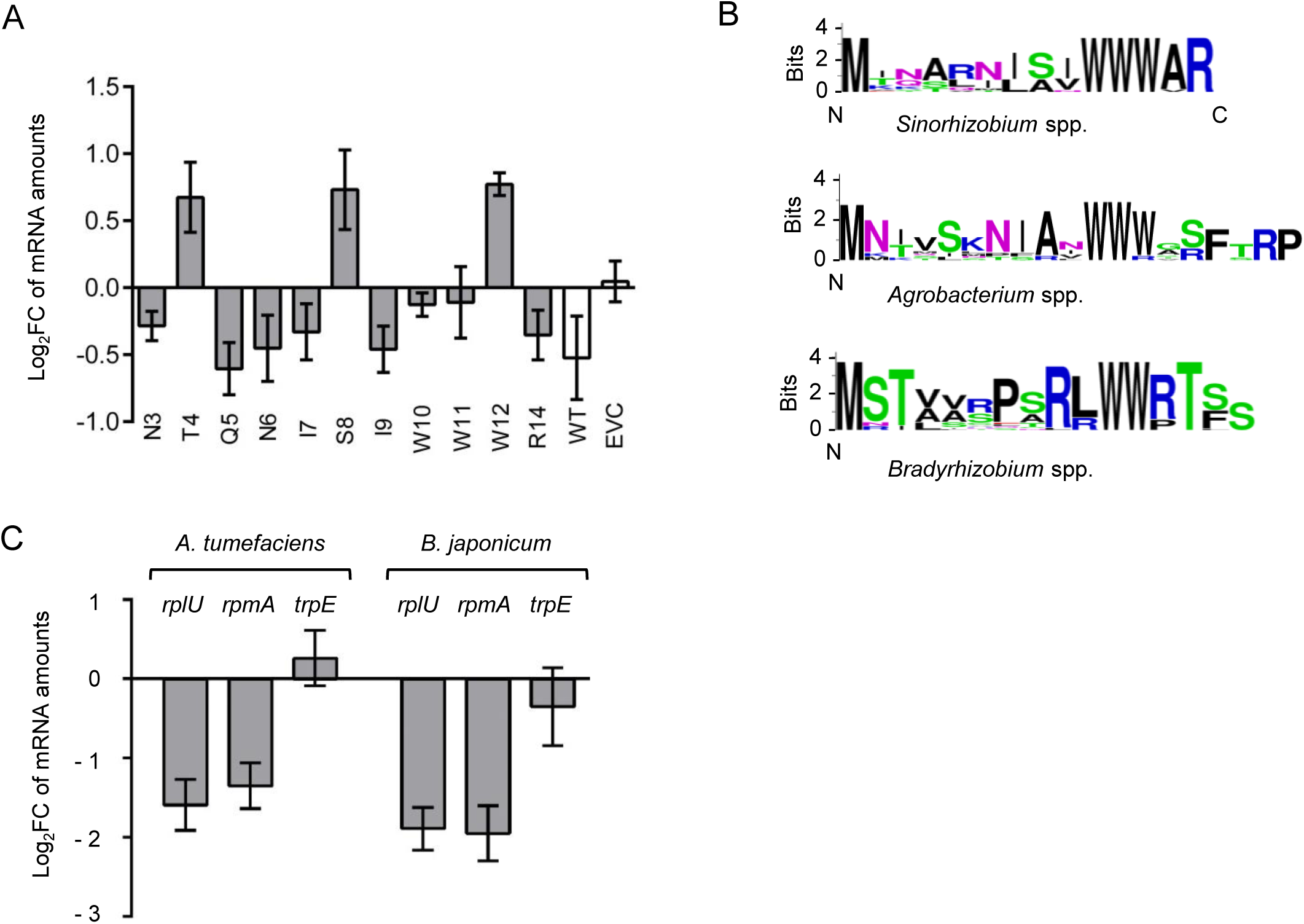
Role of peTrpL in *rplUrpmA* downregulation is conserved in Rhizobiales despite high divergence in amino acid sequence. **A)** Alanine scanning mutagenesis for analysis of functionally important residues in peTrpL of *S. meliloti*. Changes in the levels of *rpmA* were determined by qRT-PCR 10 min after addition of IPTG to induce the overproduction of peTrpL variants with the indicated aa exchanges. As EVC, *S. meliloti* 2011 cotransformed with pSRKGm and pRK4352 were used, while the strains used to overproduce wild type peTrpL (or one of its variants with aa exchanges) were cotransformed with pSRKGm-peTrpL (or one of its mutated derivatives) and pRK4352. All cultures were grown with Gm and Tc in the medium. **B)** Sequence logos for peTrpL of the *Sinorhizobium, Agrobacterium* and *Bradyrhizobium* groups (see Fig. S9). **C)** qRT-PCR analysis of the indicated mRNAs in *A. tumefaciens* and *B. japonicum* shows a decrease in the *rplUrpmA* mRNA levels upon overproduction of corresponding peTrpL homologs. Plasmid pSRKTc-Atu-peTrpL was used to induce peptide production for 10 min, and mRNA levels after induction were compared to those before induction. Due to the lack of a suitable inducible system for *B. japonicum*, Bja-peTrpL was overproduced constitutively from the chromosomally integrated, Tc-resistance-conferring plasmid pRJ-Bja-rnTrpL.

Next, we analyzed the conservation of peTrpL in the genus *Sinorhizobium* and in other *Rhizobiales*. *In silico* analysis of putative peTrpL peptides revealed several groups of conserved leader peptides that generally conform to the taxonomy (Fig. S9 and Table S2). However, no strong sequence conservation was found, with the exception of the consecutive Trp codons defining the attenuator. The consensus peTrpL sequences of the genera *Sinorhizobium, Agrobacterium* and *Bradyrhizobium* groups are shown in Fig. 12B. Surprisingly, the functionally important residues Thr4 and Ser8 of *S. meliloti* peTrpL are not conserved even within the *Sinorhizobium* group.

Despite the lack of peTrpL sequence conservation, we tested whether the role of rnTrpL and peTrpL in *rplUrpmA* regulation is conserved in *Agrobacterium tumefaciens* (which, together with *S. meliloti*, belongs to *Rhizobiaceae*), and in the more distantly related *Bradyrhizobium japonicum* (a *Bradyrhizobiaceae* member). In both species, the mRNA level of *rplUrpmA* was specifically decreased when the respective peTrpL homolog Atu-peTrpL or Bja-peTrpL was overproduced from a Tc resistance-conferring plasmid (Fig. 12C). Similar results were obtained upon overproduction of the sRNA homologs (Fig. S10). Thus, the function of peTrpL in the regulation of *rplUrpmA* seems to be conserved in these Alphaproteobacteria.

## Discussion

This proof of principle study establishes that bacterial leader peptides can exert conserved functions in *trans*. Our data provide strong evidence that, in *S. meliloti*, the 14-aa leader peptide peTrpL is involved in destabilization of *rplUrpmA* mRNA upon exposure to representative translation-inhibiting antibiotics. This downregulation of *rplUrpmA* expression by peTrpL depended on the presence of the attenuator sRNA rnTrpL and Tc or other translation-inhibiting antibiotics (Fig. 1G, Fig. 2, Fig. 3 and Fig. 11A).

We provide conclusive evidence that the sRNA rnTrpL base pairs with *rplU* and that this direct interaction downregulates the *rplUrpmA* mRNA levels (Fig. 4). The identification of *rplUrpmA* as a direct target of rnTrpL extends the list of confirmed interactions of this particular sRNA. Interestingly, the downregulation of the previously identified target *trpDC* (Melior et al., 2019) did not depend on peTrpL or Tc (Fig. 1C and Fig. S2), suggesting that interactions of rnTrpL with its diverse targets may be facilitated by different mechanisms. The special requirement of both peTrpL and an antibiotic for *rplUrpmA* downregulation probably ensures that rnTrpL can be efficiently redirected from other mRNAs (such as *trpDC*) to this alternative target under conditions of translation inhibition. In support of this hypothesis, we were able to demonstrate *in vitro* the redirection of the rnTrpL sRNA from *trpD* to its Tc-dependent target *rplU* (Fig. 8D).

The antibiotic-triggered, posttranscriptional downregulation of *rplUrpmA* identified in this study probably serves to adjust the production and/or function(s) of ribosomes. Lower levels of *rplUrpmA* mRNA may negatively influence ribosome biogenesis and/or result in ribosomes lacking the L21 and L27 proteins. While the function of L21 is not clear, *E. coli* L27-deficient mutants were shown to have reduced peptidyl transferase activities (Maguire et al., 2005). Interestingly, a two-fold reduction of *rplUrpmA* expression in *Pseudomonas aeruginosa* leads to an increased expression of multidrug efflux pump genes and increased resistance to aminoglycosides. This was explained by attenuation of transcription termination, caused by pausing of L21- and L27-less ribosomes (Lau et al., 2012). It therefore seems reasonable to suggest that, in *S. meliloti*, the downregulation of *rplUrpmA* by rnTrpL and peTrpL may have similar adaptation effects mediated by specific changes in translation.

Another notable finding of our work is the generation of the sRNA rnTrpL upon translation inhibition, even under conditions of Trp insufficiency. We show that bacteria utilize the same uORF and two mutually exclusive RNA structures for two different purposes: 1) to regulate *trpE(G)* expression by transcription attenuation in response to Trp availability (Bae and Crawford, 1990) and 2) to achieve transcription termination between *trpL* and *trpE(G)* upon translation inhibition even under conditions of Trp shortage (Fig. 10A). In most ribosome-dependent transcription attenuators of amino acid biosynthesis operons, the relevant codons are located in the 3′-half of the uORF (Vitreschak et al., 2004). Translation inhibition at such attenuators would also generate sRNAs with potential functions in *trans*. The rnTrpL sRNA production upon ribosome pausing in the 5′-part of the uORF *trpL* may have additional implications. We note that, in all *Sinorhizobium* species, two asparagine and one glutamine codon(s) are located in the first half of *trpL*. Pausing of the ribosome at these codons, for example upon shortage of aminoacylated tRNA^Asn^ and tRNA^Gln^, could also result in rnTrpL production. Thus, *trpL* is well-suited for sensing nitrogen depletion in *Sinorhizobium*. Furthermore, since all biosynthetic, ribosome-dependent attenuators have the capacity to respond to translation inhibition, it seems reasonable to suggest that attenuator sRNAs are generated under stringent response conditions and may play important roles in the stress response and/or subsequent recovery pathways.

We note that the *trp* attenuator of the 5′-leader of *trpE(G)* is strikingly multifunctional (Fig. 10C, Fig. 13). Few other bacterial 5′-UTRs are known to respond to different intercellular signals or to act both in *cis* and in *trans*. The 5′-UTR of *mgtA,* which encodes a Mg^2+^ transporter in *Salmonella,* is one of the few examples of a *cis*-acting attenuator that is capable of sensing disparate signals. It harbors a proline-rich uORF and can respond to proline shortage or a decline in Mg^2+^ concentrations (Park et al., 2010; Chadani et al., 2017). Metabolite-binding riboswitches that can act in *cis* and in *trans* were described in *Listeria* and *Enterococcus* (Loh et al., 2009; DebRoy et al., 2014; Mellin et al., 2014). The *trp* attenuator of *S. meliloti* combines similar mechanisms in an exceptional complexity: 1) in *cis*, it senses two disparate signals, Trp availability (Bae and Crawford, 1990) and translation inhibition (this work); 2) the attenuator sRNA, which is released upon transcription termination, is a *trans*-acting sRNA that regulates several genes including *sinI* (Baumgardt et al., 2016), the *trpDC* operon (Melior et al., 2019) and *rplUrpmA* (this work); 3) the encoded leader peptide peTrpL is also functional in *trans* and (together with rnTrpL), can mediate the antibiotic-dependent *rplUrpmA* downregulation.

**Figure 13.**
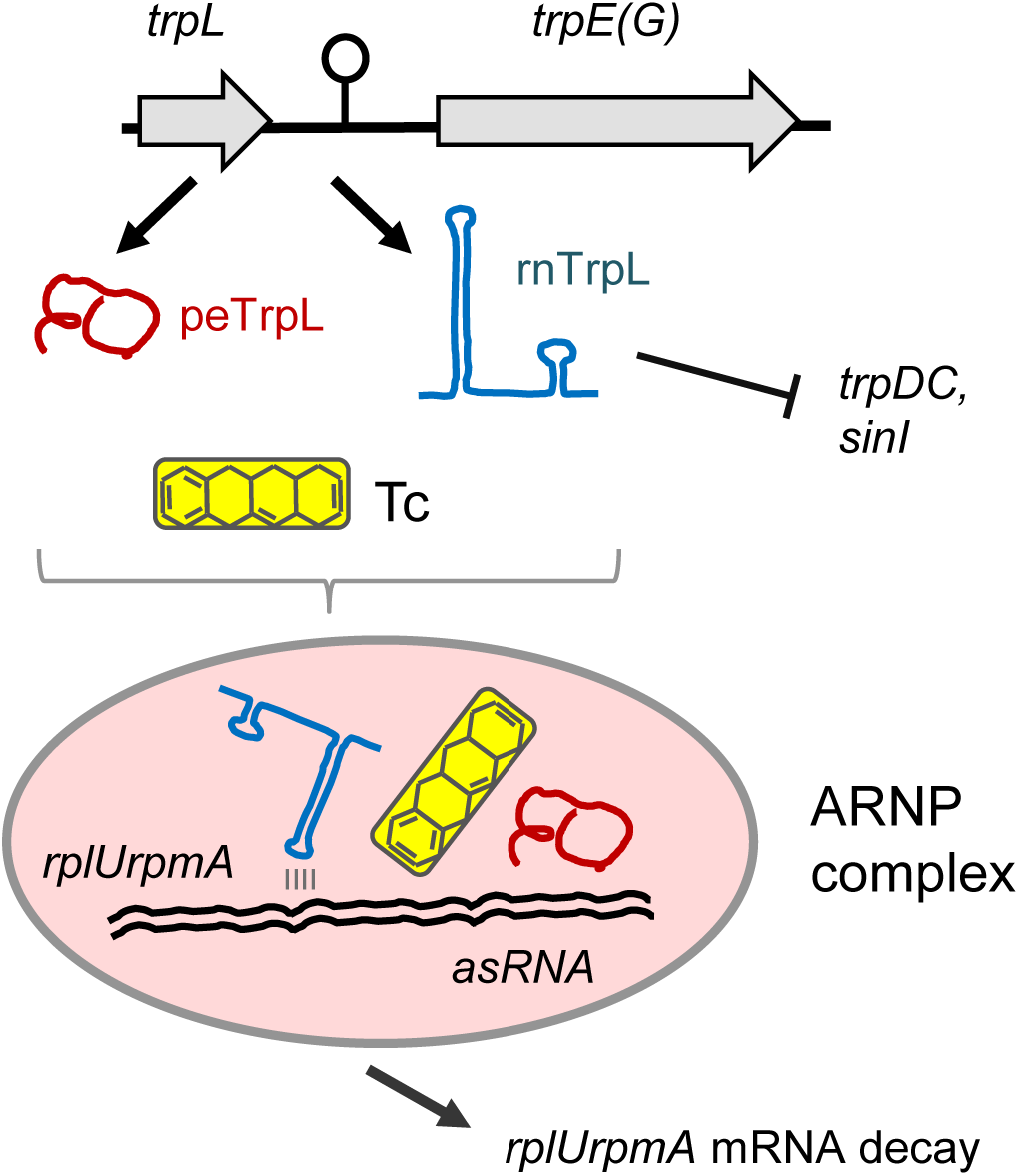
Model of the peTrpL leader peptide function in ARNP formation and *rplUrpmA* destabilization upon exposure of *S. meliloti* to tetracycline. The leader peptide and the antibiotic redirect the attenuator sRNA rnTrpL from antibiotic-independent targets such as *trpDC* to *rplUrpmA* mRNA. The asRNA anti-*rplUrpmA*, which is induced upon exposure to Tc and increases the ARNP complex formation, is shown in a duplex with *rplUrpmA* mRNA. Base-pairing between rnTrpL and a specific *rplU* sequence is necessary for *rplUrpmA* destabilization.

Interestingly, the posttranscriptional regulation of *rplUrpmA* by peTrpL and rnTrpL is intimately linked to the presence of translation-inhibiting antibiotics. The levels of peTrpL in strain 2011 were strongly increased upon exposure of the cells to Tc, in line with a specific function of the leader peptide under this condition (Fig. 10B). Further, *trpL* was also shown to be necessary for *rplUrpmA* decrease in response to the translation inhibitors Tc, Em, Cl and Km, but not to Rf, which inhibits transcription (Fig. 11A). Moreover, each of these translation-inhibiting antibiotics, but not Rf, supported the production of ARNP complexes (Fig. 11B). The observed complex disassembly at low Tc concentrations (or during wash steps with buffer lacking antibiotics) suggests that the antibiotics participate in allosteric interactions in the ARNPs. Such interactions would allow for fast and flexible complex formation (leading to *rplUrpmA* destabilization) or disassembly (abrogating the *rplUrpmA* destabilization) in response to changes in the intracellular antibiotics concentration. Such fast, posttranscriptional regulation would be especially useful for *S. meliloti* in its soil and rhizosphere habitat, where many antibiotic-producing microorganisms are present and competition for nutrients and space is high (Naamala et al., 2016). Therefore, we suggest that the antibiotic-triggered *rplUrpmA* downregulation by peTrpL is a part of an adaptation mechanism.

Unexpectedly, asRNAs were present in the ARNPs purified from *S. meliloti* (Fig. 6, Fig. 11B). However, these asRNAs did not seem to be absolutely necessary for *rplUrpmA* downregulation by rnTrpL *in vivo*, because the asRNAs were not co-induced in the two-plasmid assay showing the base-pairing between rnTrpL and *rplU* (Fig. 4). Further, asRNAs were dispensable for the reconstitution of core ARNPs *in vitro* (Fig. 9A). However, our data suggest that the anti-*rplUrpmA* RNA, which is specifically induced under antibiotic stress, may increase the efficiency of ARNP formation (Fig. 9B). Thus, *in vivo* the asRNAs may support the redirection of rnTrpL from *trpDC* to *rplUrpmA* upon antibiotic exposure.

The mechanism reported here for *rplUrpmA* destabilization adds another facet to the molecular responses of bacteria to antibiotics. It is well known that antibiotics are used as allosteric effectors of TetR-type repressors to induce the transcription of resistance genes (Hillen and Berens, 1994; Reichheld et al., 2009). Additionally, ribosome stalling due to translation-inhibiting antibiotics has regulatory effects on transcription attenuators. Ribosomes stalled at uORFs can either prevent transcription termination and thus induce the expression of downstream resistance genes (Dar et al., 2016) or, as described here, the stalled ribosomes promote transcription termination to generate a *trans*-acting sRNA (Fig. 9A). Moreover, as shown here, bacteria can directly use the antibiotics for ARNP formation in the process of mRNA destabilization.

Surprisingly, despite the lack of sequence conservation beyond the Trp residues, the function of peTrpL in *rplUrpmA* downregulation is conserved among soil Alphaproteobacteria, such as the plant pathogen *A. tumefaciens* and the soybean symbiont *B. japonicum*. This probably reflects molecular adaptation to (or co-evolution with) RNA interaction partners in the respective host. The functional conservation of peTrpL suggests that ARNP-based regulatory mechanisms may also be conserved in other bacteria.

In summary, our work establishes a role of the leader peptide peTrpL in *trans*, extends the target spectrum of the attenuator sRNA rnTrpL, indicates that the *trp* attenuator can sense translation inhibition as a second signal, and demonstrates the formation of antibiotic-supported ribonucleoprotein complexes. We are convinced that the data described here will inspire future research on multifunctional 5′-UTRs and in particular on additional functions of *trans*-acting leader peptides. More generally, our data encourage future analyses of novel interactions between antibiotics, nucleic acids and proteins and of the underlying mechanisms that control and regulate these interactions.

## Material and Methods

### Cultivation of bacteria, conjugation, IPTG induction, antibiotics exposure and fluorescence measurement

*E. coli* was cultivated in LB, *S. meliloti* 2011 and its derivatives in rich TY or minimal GMX medium, *A. tumefaciens* NTL4 in TY, and *B. japonicum* 110spc4 in in PSY medium. Growth conditions and selective antibiotic concentrations were as previously described (Melior et al., 2019; Mesa et al., 2008). Alphaproteobacteria were cultivated to an OD_600nm_ of 0.5, and then processed. Cloning procedures were performed essentially as described by Sambrook et al. (1989).

All oligonucleotides (primers) are listed in Table S3. They were synthesized by Microsynth (Balgach, Switzerland). The used plasmids are listed in Table S4. As vectors the conjugative, broad-host range vectors pRK4352 (Mank et al., 2012), pSRKGm or pSRKTc (Khan et al., 2008), which can replicate in *S. meliloti* and *A. tumefaciens*, were used as described (Melior et al., 2019). Alternatively, plasmids pRJPaph-MCS (Hahn et al., 2016) or a derivative of pSUP202pol4 (Fischer et al., 1993) containing 300 nt of the 3′ *exoP* region (Schlüter et al., 2015) were used as vectors for chromosomal integration *B. japonicum* or *S. meliloti*, respectively. Inserts were amplified by PCR or, for very short inserts (e.g. *trpL* ORFs with codons exchanged for synonymous codons, or with mutated codons), complementary oligonucleotides were annealed and cloned. pLK64 was used as an *egfp* template (McIntosh et al., 2008). Plasmids were transferred from *E. coli* S17-1 by conjugation (Simon et al., 1983). Constitutively overproducing strains were tested immediately after conjugation. IPTG was used at a final concentration of 1 mM for the indicated periods of time.

For short-term exposure of *S. meliloti* strains harboring a Tc-resistance plasmid, 20 µg/ml Tc was used. Further, following subinhibitory antibiotic concentrations were used for strains lacking resistance plasmids: 1.5 µg/ml Tc, 27 µg/ml Em, 9 µg/ml Cl, 3 µg/ml Rf, 45 µg/ml Km. A control culture was exposed in parallel to the respective solvent (ethanol, methanol or water). EGFP fluorescence was measured in Tecan Infinite M200 reader 20 min after IPTG addition to cultures of strains containing two plasmids (pSRKGm- and pSRKTc-constructs for IPTG-induced expression of *egfp* reporter fusions and the sRNA, respectively).

For more details, see the Supplementary Methods File.

### RNA methods

Total RNA of *S. meliloti* and *A. tumefaciens* was isolated using TRIzol and hot-phenol (for Northern blot hybridization and qRT-PCR analysis) or using RNeasy columns (for RNA half-live measurements; Qiagen, Hilden, Germany) as described (Melior et al., 2019). Northern Blot hybridization with radioactively labeled oligonucleotide probes was also previously described (Melior et al., 2019). Strand-specific, real time, quantitative reverse transcriptase PCR (qRT-PCR) was performed with the Brilliant III Ultra Fast SYBR® Green QRT-PCR Mastermix (Agilent, Waldbronn, Germany): 5 µl Master Mix (supplied), 0.1 µl DTT (100 mM; supplied), 0.5 µl Ribo-Block solution (supplied), 0.4 µl water, 1 µl of the reverse primer (10 pmol/µl), and 2 µl RNA (20 ng/µl) were assembled in a 9-µl reaction mixture. After cDNA synthesis, the samples were incubated for 10 min at 96 °C to inactivate the reverse transcriptase.

After cooling to 4 °C, 1 µl of the second primer (10 pmol) was added, and real-time PCR was performed starting with 5 min incubation at 96 °C. Used primers and their efficiencies (determined by PCR of serial two-fold RNA dilutions) are listed in Table S3. A spectrofluorometric thermal cycler (Biorad, München, Germany) was used and the quantification cycle (Cq), was set according to (Bustin et al., 2009). As a reference, *rpoB* mRNA (for steady state level analyses) or 16S rRNA (for half-lives determination) was used (Baumgardt et al., 2016; Melior et al., 2019). Stability of mRNA was determined after addition of rifampicin was added to a final concentration of 600 μg/ml as described (Melior et al., 2019). The Pfaffl formula was used to calculate fold changes of mRNA amounts (Pfaffl, 2001). For qRT-PCR analysis of total RNA, the analysis of the gene of interest (e.g. *rpmA*) and of the reference gene *rpoB* were performed using portions of the same DNA-free RNA sample, and log_2_fold changes of mRNA levels after induction by IPTG and/or exposure to antibiotics were determined. For qRT-PCR analysis of coimmunoprecipitated RNA (CoIP-RNA) or RNA co-purified with MS2-rnTrpL, the real-time RT-PCR of the gene of interest was performed using a CoIP-RNA sample, while total RNA of the same culture (harvested prior to cell lysis for CoIP) was used for the *rpoB* real-time RT-PCR. Then, the Pfaffl formula was used to calculate the fold enrichment of specific RNAs by CoIP with 3×FLAG-peTrpL or by affinity chromatography with MS2-MBP, in comparison to the corresponding mock purifications.

RNA-seq was performed by Vertis Biotechnologie AG (Freising, Germany). cDNA reads were mapped as described (Sharma et al., 2010).

For *in vitro* transcription the MEGAshortscript T7 kit (Ambion) was used according to the manufacturer instructions. The T7 promoter sequence was either integrated in one of the primers for PCR amplification of the template, or it was present in oligonucleotides that were annealed to obtain double-strand template (see Table S3). TURBO-DNase was used to remove the DNA template (The *in vitro* transcript was extracted with acidic phenol, precipitated with ethanol and analysed in a 10% polyacrylamide-urea gel after staining with ethidium bromide.

For more details, see the Supplementary Methods File.

### Coimmunoprecipitation of RNA with 3×FLAG-peTrpL and MS2-rnTrpL affinity purification

*S. meliloti* 2011 strains containing the plasmid pSRKGm-3×FLAG-peTrpL were used for the coimmunoprecipitation (CoIP) 10 min after addition of IPTG. For the mock CoIP control pSRKGm-peTrpL was used instead. Cell pellets were resuspended in 5 ml buffer B (20 mM Tris, pH 7.5, 150 mM KCl, 1 mM MgCl_2_, 1 mM DTT) containing 10 mg/ml lysozyme, 2 µg/ml Tc and 1 tablet of protease inhibitor cocktail (Sigma Aldrich) per 40 ml buffer. Cells were lysed by sonication and 40 µl Anti FLAG® M2 Magnetic Beads (Sigma Aldrich) were added to the cleared lysate. After incubation at 4 °C for 2 h, the beads were collected and split into two portions. One sample was washed 3 times with 500 µl buffer B containing an antibiotic (at the indicated concentration), while the second sample was washed with buffer alone, and the beads were resuspended in 50 µl buffer B. Coimmunoprecipitated RNA was purified using TRIzol. MS2-rnTrpL affinity purification from *S. meliloti* 2011 strains containing the plasmid pRK-MS2-rnTrpL was performed using MS2-MBP fusion protein bound to amylose beads (New England Biolabs) as described by Smirnov et al. (2016). The rnTrpL derivative was tagged at the 5′-end. A mock purification using bacteria containing plasmid pRK-MS2 was performed. As described above, the beads were washed with a buffer that did or did not contain an antibiotic. The RNA and protein content of the elution fractions was analyzed. For more details, see the Supplementary Methods File.

### Tricine-SDS-PAGE and Western blot analysis

Tricine-SDS-PAGE was conducted according to Schägger (2006). The 16 % polyacrylamide separating gel (acrylamide:bisacrylamide 19:1) contained 8 % glycerol. Detection of FLAG-tagged proteins transferred onto a PVDF membrane was performed with monoclonal anti-FLAG M2-peroxidase (HRP) antibodies (Sigma-Aldrich) and Lumi-Light Western Blotting Substrate Kit (Roche). FLAG-tagged FLAG-FimV Protein (Rossmann et al., 2019) was used as a positive control.

### Mass spectrometry

For identification of proteins in a gel slice stained with Coomassie Brilliant Blue, the band was destained and digested with trypsin as reported elsewhere (Bonn et al., 2014). To recover the peptides, gel pieces were covered with ultra-pure water and incubated 15 minutes in an ultrasonic water bath. Peptides derived from in-gel digestion were loaded on an EASY-nLC II system (Thermo Fisher Scientific) equipped with an in-house built 20-cm column (inner diameter 100 mm, outer diameter 360 mm) filled with ReproSil-Pur 120 C18-AQ reversed-phase material (3 mm particles, Dr. Maisch GmbH). Elution of peptides was executed with a nonlinear 80-min gradient from 1 to 99% (v/v) solvent B (0.1% (v/v) acetic acid in acetonitrile) with a flow rate of 300 nl/min and injected online into a LTQ Orbitrap XL (Thermo Fisher Scientific). The survey scan at a resolution of R = 30.000 and 1 x 10^6^ automatic gain control target in the Orbitrap with activated lock mass correction was followed by selection of the five most abundant precursor ions for fragmentation. Singly charged ions as well as ions without detected charge states were excluded from MS/MS analysis.

For quantification of peTrpL abundance, protein extracts were diluted in 50 mM TEAB-Buffer, pH 8.0 (Sigma-Aldrich) to a final concentration of 0.5 µg/µl. After protein reduction (2.5 mM TCEP, Tris-(2-carboxyethyl) phosphine hydrochloride, Invitrogen) at 65 °C for 45 min, thiols were alkylated in 5 mM iodacetamide (Sigma) for 15 min at 25 °C in the dark. For protein digestion, trypsin (Promega) was added in an enzyme-to-substrate ratio of 1:100. After 14 h at 37 °C, digestion was terminated by adding concentrated HCl to a final concentration of 600 mM and peptides were purified by C18 Zip tips (Pierce). Prior measurement samples were spiked with synthetic peptides containing an isotopically labeled amino acid (JPT Peptide Technologies) to a final concentration of 50 fmol/µl. The peptides derived from in-solution digests were loaded on an EASY-nLC 1000 system (Thermo Fisher Scientific) equipped with an in-house built 20-cm column (see above). Elution of peptides was executed with a nonlinear gradient from 1 to 99% (v/v) solvent B (0.1% (v/v) acetic acid in acetonitrile) with a flow rate of 300 nl/min and injected online into a TSQ Vantage (Thermo Fisher Scientific). The selectivity for both Q1 and Q3 were set to 0.7 Da (FWHM). The instrument was operated in SRM mode applying a collision gas pressure of 1.2 mTorr in Q2. The collision energy was optimized before analysis. All monitored transitions and the optimized collision energy can be found in Table S5.

### Processing of mass spectrometry data

For identification of peptides from MS-spectra, a database search was performed with Sorcerer-SEQUEST (4.0.4 build, Sage-N Research) using the Sequest algorithm against a target-decoy integrated proteogenomic database (iPtgxDB; https://iptgxdb.expasy.org/), which also contained sequences of common laboratory contaminants and FLAG-tagged peTrpL (total entries: 320482). The *S. meliloti* 2011 iPtgxDB was created by integrating and consolidating the annotations of the chromosome (NC_020528) and two plasmids (NC_020527 and NC_020560) from RefSeq (Tatusova et al., 2016) and Genoscope (Vallenet et al., 2013), with predictions from Prodigal (Hyatt et al., 2010), ChemGenome (Singhal et al., 2008) and a modified form of six-frame predicted ORFs (Omasits et al., 2017). The database search was based on a strict trypsin digestion with two missed cleavages permitted. Oxidation of methionine and carbamidomethylation of cysteine were considered as variable modifications. The mass tolerance for precursor ions was set to 10 ppm and the mass tolerance for fragment ions to 0.5 Da. Validation of MS/MS-based peptide and protein identification was performed with Scaffold V4.7.5 (Proteome Software, Portland, USA), and peptide identifications were accepted if they exceeded the following thresholds: deltaCn greater than 0.1 and XCorr scores greater than 2.2, 3.3 and 3.75 for doubly, triply and all higher charged peptides, respectively. Protein identifications were accepted if at least 2 identified peptides were detected for proteins with a molecular weight of 15 kDa and higher. For proteins smaller than 15 kDa the identification of one unique peptide fulfilling the criteria mentioned above was sufficient for an identification. Normalized spectrum abundance factors (Zybailov et al., 2006) were used as proxy for protein abundance in the sample.

All raw files from targeted MS were processed using Skyline 4.2 (MacLean et al., 2010). A peptide ratio of native and heavy species was based on five transitions. Peptide ratios of three biological replicates were weighted according to their Dot product before being averaged. The concentration of peTrpL in the sample was calculated based on the peptide ratios and the added amount of heavy peptide.

### Complex disassembly, reassembly and reconstitution

ARNP complex was eluted from the MS2-MBP-amylose beads in buffer C (20 mM Tris, pH 8.0, 150 mM KCl, 1 mM MgCl_2_, 1 mM DTT) containing 2 µg/ml Tc (High-Tc conditions). For disassembly and reassembly in the presence of synthetic 3×FLAG-peTrpL (Thermo Fischer Scientific), 5 µl peptide solution (1 µg/µl; 222 pmol/µl) was added to 150 µl of the ARNP sample and three 50-µl aliquots of the sample were processed in parallel. One control aliquot was diluted five-fold with Tc-containing buffer C and incubated for 10 min under High-Tc conditions (no complex disassembly). The second aliquot was diluted five-fold in buffer C lacking Tc, incubated at the final Tc concentration of 0.4 µg/ml (Low-Tc conditions; complex disassembly), and then 4 µl Tc solution (100 µg/ml, dissolved in ethanol) was added to increase the Tc concentration of the diluted sample to 2 µg/ml. Subsequently the sample was incubated for complex reassembly at the High-Tc conditions for 7 min. The third aliquot was diluted five-fold in buffer C lacking Tc and incubated for 3 min before 4 µl ethanol was added and the sample was incubated for additional 7 min under the Low-Tc conditions. Then 70 µl MS2-MBP-amylose beads in buffer C was added to each of the three samples, and affinity chromatography was performed. Alternatively, CoIP was performed with 20 µl magnetic beads with anti-FLAG antibodies. RNA and protein content of the elution samples were analyzed.

Reconstitution reactions with entirely synthetic components were performed using MS2-rnTrpL (100 ng; 2 pmol), mini-*rplU* (100 ng; 4.4 pmol), peTrpL (50 ng, 27 pmol) and 3×FLAG-peTrpL (50 ng, 11 pmol) in buffer C, in a final volume of 50 µl. Addition of vitro transcribed antisense RNA (100 ng, 4.4 pmol) that is complementary to the mini-*rplU* transcript, unrelated, control *in vitro* transcript *crtA* corresponding to a *Rhodobacter sphaeroides* sequence (100 ng or 300 ng), control yeast tRNA (Sigma) (100 ng or 300 ng) and/or of Tc (to a final concentration of 2 µg/ml) is indicated. The samples were incubated for 20 min at 20 °C, and then 10 µl of MS2-MBP-amylose beads were used to purify reconstituted, MS2-rnTrpL-containing ARNP. Alternatively, reconstituted, 3×FLAG-peTrpL-containing complexes were isolated by CoIP with anti-FLAG antibodies. RNA was purified from the elution fractions and was analyzed by Northern blot hybridization and/or qRT-PCR. For other details, see text and figure legends.

### Computational analysis

281 genomes of Rhizobiales were downloaded from GenBank (Benson et al., 2013) in August 2018. Intergenic regions of length 300 upstream of the gene *trpE* were extracted. We then considered all open reading frames containing at least two consecutive UGG codons and converted them into amino acid sequences of candidate leader peptides. The peptides were aligned using MUSCLE with default parameters (local alignment, open gap penalty –10, extend gap penalty –1) (Edgar, 2004). Hanging N-termini were trimmed manually, retaining the condition that candidate leader peptides should start with methionine. Phylogenetic trees were constructed based on the constructed distance matrix using R package “cluster” with default parameters (Maechler et al., 2018) by the Ward2 method (Murtagh and Legendre, 2014) and visualized using R package “ggplot2” (Wickham, 2016). Sequence logos were constructed using WebLogo (Crooks et al., 2004). The validity of predicted leader peptides was assessed by construction of alternative RNA secondary structures characteristic of attenuators using ad hoc Python scripts for the identification of overlapping RNA helices.

RNA-RNA interactions were predicted by IntaRNA (Mann et al., 2017) as described (Melior et al., 2019). Primer for qRT-PCR analysis were designed using Primer3 (Untergasser et al., 2012). Secondary structures of mini-*rplU* and mini-*trpD* were predicted by MFOLD (Zuker, 2003).

## Data availability

The RNA-Seq and RIP-Seq data discussed in this publication have been deposited in NCBI’s Gene Expression Omnibus (Edgar et al., 2002); accession number GSE118689. The MS data discussed in this publication have been deposited to the ProteomeXchange Consortium via the PRIDE partner repository (Vizcaíno, J. A. et al., 2016) with the dataset identifier PXD 015368 (Reviewer account details: Username, Password: O64HhpDz). The *S. meliloti* 2011 iPtgxDB is available at https://iptgxdb.expasy.org/database/.

## Acknowledgments

We are grateful to Jürgen Bartel (University of Greifswald, Germany) for excellent technical assistance in mass spectrometry. We thank Janina Gerber and Oliver Puckelwaldt (University of Giessen, Germany) for help in some experiments. The peTrpL computational analysis in Alphaproteobacteria was initiated at the Summer School of Molecular and Theoretical Biology supported by the Zimin Foundation. We are grateful to Jörg Vogel (University of Würzburg, Germany) for sending us the *E. coli* strain for MS2-MBP purification and for providing the protocols for MS2-MBP affinity chromatography.

## Author contributions

Conceptualization, E.EH., H.M.; Methodology, E.E.H., H.M., S.M., Z.C.; Investigation, H.M., S.M., M.S., S.L., R.S., S.A., A.S., S.B.W., K.B.; Formal analysis, H.M., S.M., S.L., M.S., K.U., A.R.V., A.S., Z.C., C.H.A.; Writing – Original Draft, E.E.H., H.M., J.Z.; Writing – Review and Editing, E.E.H., H.M., S.M., Z.C., J.Z., C.H.A.; Visualization, E.E.H., H.M., S.M., S.L., A.S., Z.C.; Supervision, E.E.H., Z.C., D.B., C.H.A.; Funding Acquisition, E.E.H.

## Funding

This work was funded by DFG (Ev 42/6-1; BE 3869/5-1 and Ev 42/7-1 in SPP2002; GRK2355 project number 325443116). S.L. was supported by the China Scholarship Council (No. 201708080082); Z.C. was supported by the Russian Science Foundation (18-14-00358); J.Z. was supported by the DFG (SFB1021, A01 and B01). A.R.V. was supported by a grant from the Swiss National Science Foundation (156320).

## Declaration of interests

The authors declare no competing interests.

## Supplementary Files

Supplementary Methods

Supplementary Figures

Table S1, Mass spectrometry results for the 13 kDa band co-purified with MS2-rnTrpL.

Table S2, List of putative peTrpL sequences of Rhizobiales members.

Table S3, Oligonucleotides used in this work.

Table S4, Plasmids used in this work.

Table S5, Mass spectrometry data for the quantified peTrpL peptide in both isotope forms.

