## Supplementary Figures 1 to 10 for "The bacterial leader peptide peTrpL has a conserved function in antibiotic-dependent posttranscriptional regulation of ribosomal genes"

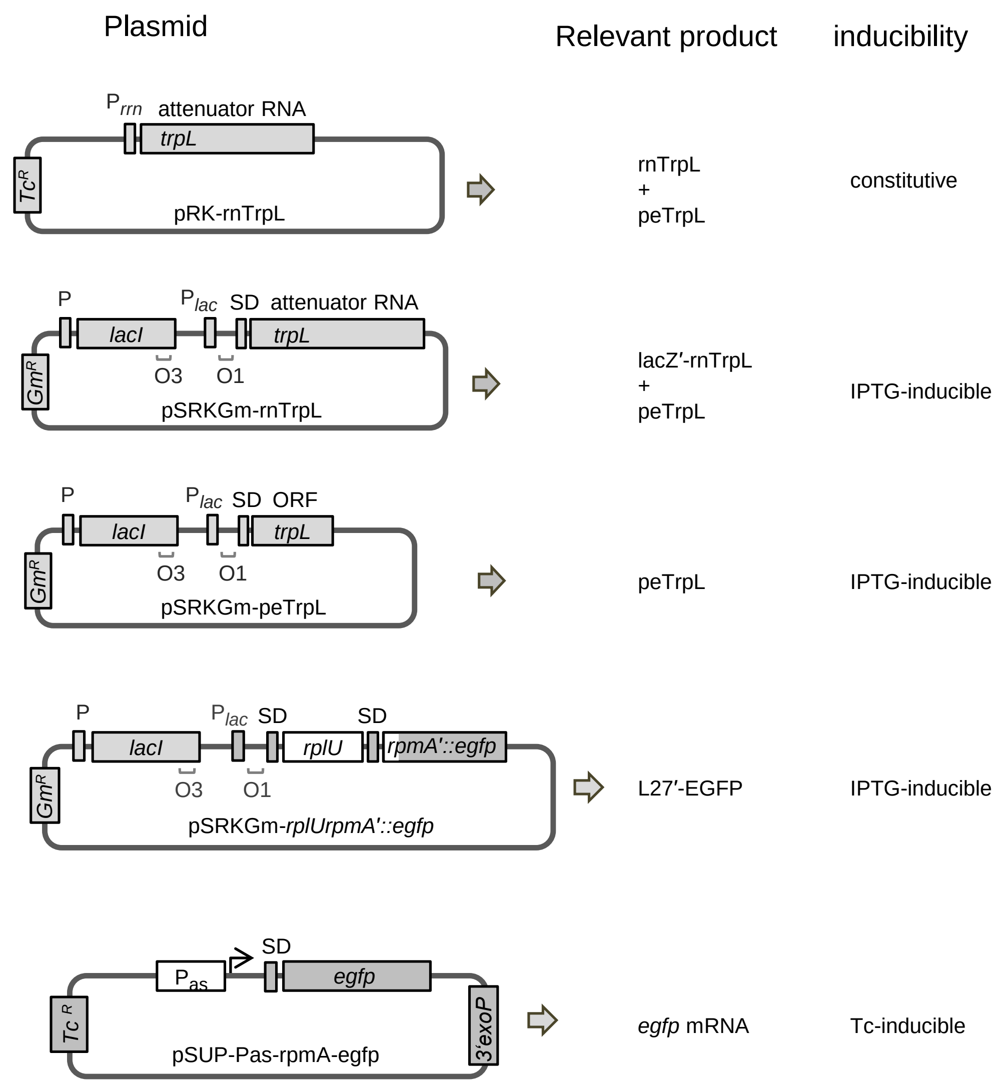


**Figure S1. Plasmids for constitutive and induced overproduction of rnTrpL, lacZ′-rnTrpL, and reporter protein or mRNA, respectively, with indicated products and inducibility.** In addition to the here shown pSRKGm plasmids with a Gm resistance gene, similar pSRKTc-constructs with a Tc resistance gene were also used.

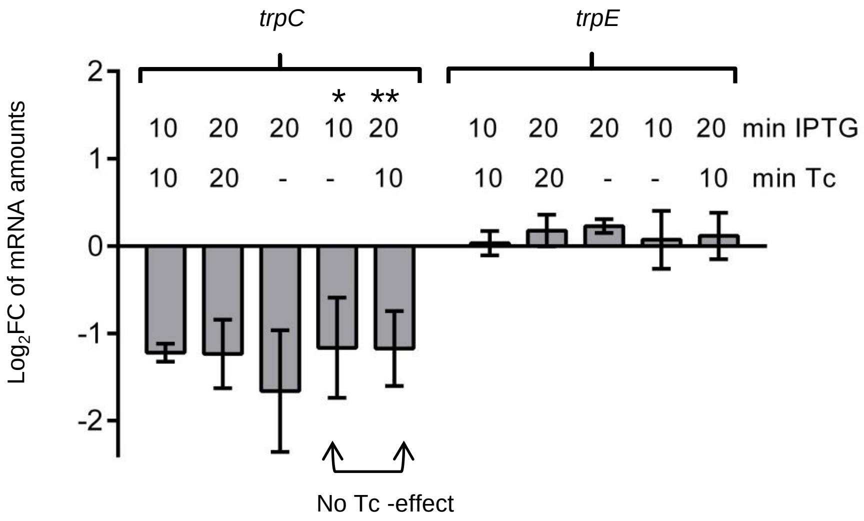


**Figure S2. Regulation of *trpC* by rnTrpL does not depend on Tc.** Effects of Tc and rnTrpL on *trpC* expression in strain 2011Δ*trpL* (pSRKGm-rnTrpL, pRK4352) were analyzed. The culture was grown first in medium with Gm and Tc. Then the cells were washed and grown for 4 h without Tc. IPTG was added at time point 0 min and 10 min later (time point 10 min, *), Tc was also added. 10 min after Tc addition (corresponds to 20 min after IPTG addition; time point 20 min, **), the experiment was completed. RNA was isolated and the *trpC* levels at the time points 10 and 20 min were compared to the level at the time point 0 by qRT-PCR (see also Fig. 3A). Suitable controls were conducted. Shown are means and standard deviations from three independent experiments, each performed in duplicates. As a negative control (gene not regulated by rnTrpL in *trans*), *trpE* was used.

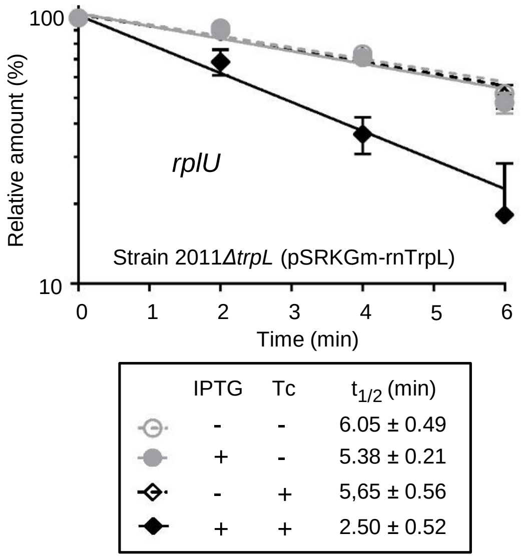


**Figure S3. Determination of *rplU* mRNA stability in strain 2011*ΔtrpL* (pSRKGm-rnTrpL) upon lacZ′-rnTrpL induction and/or Tc exposure.** Bacterial cultures were grown with Gm in the medium. A culture was split in four portions, which were treated differently in respect to IPTG and Tc addition (indicated). 10 min after induction of rnTrpL transcription by IPTG and/or Tc addition (1.5 µg/ml), rifampicin (Rf) was added to stop cellular transcription. In parallel, non-induced cultures that were exposed or not exposed to Tc, were also treated with Rf. Relative mRNA levels were determined, the mRNA level at time point 0 (before Rf addition) was set to 100%, the relative mRNA level values were plotted against the time, and the mRNA half-lives were calculated. Shown are the results from three independent transcription inhibition experiments. The qRT-PCRs reactions were performed in technical duplicates (means with standard deviations are indicated).

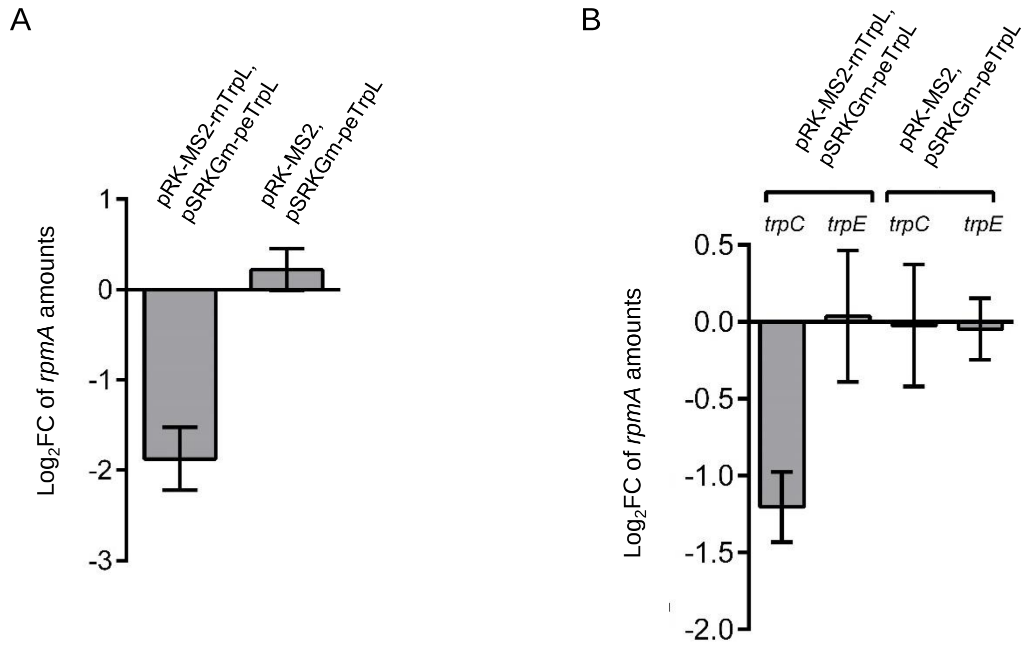


**Figure S4. Functionality of the constitutively produced, tagged sRNA MS2-rnTrpL. A)** The tagged sRNA mediates down-regulation of *rpmA* upon peTrpL induction. Strain 2011*ΔtrpL* (pRK-MS2-rnTrpL, pSRKGm-peTrpL) was grown in medium with Gm and Tc. IPTG was added to induce peTrpL production. The *rpmA* mRNA level 10 min post induction was compared to the level before induction by qRT-PCR. A decrease in the *rpmA* level was not detected in the control strain 2011*ΔtrpL* (pRK-MS2, pSRKGm-peTrpL), which lacks the sRNA. **B)** The tagged sRNA down-regulates *trpC*. The levels of the mRNAs *trpC* (peTrpL-independent target of rnTrpL) and *trpE* (negative control) of strains 2011*ΔtrpL* (pRK-MS2-rnTrpL, pSRKGm-peTrpL) and 2011*ΔtrpL* (pRK-MS2, pSRKGm-peTrpL) were compared to the levels in the EVC 2011*ΔtrpL* (pRK4352, pSRKGm). IPTG was not added. Shown are means and standard deviations from three independent experiments, each performed in duplicates. The used plasmids are indicated.

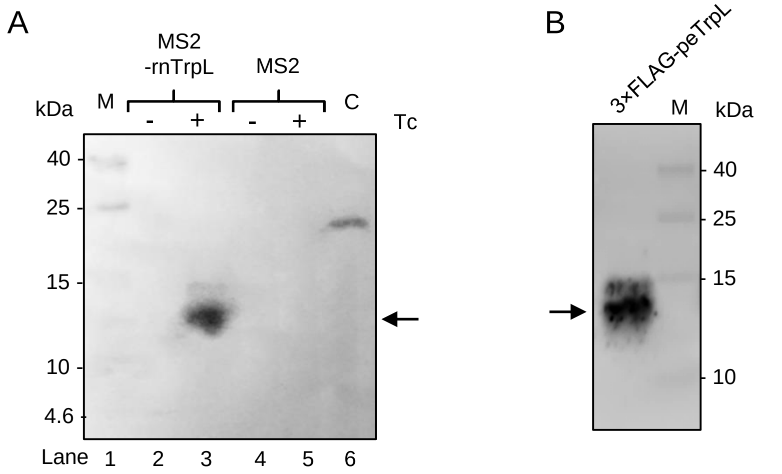


**Figure S5. Western blot detection of 3×FLAG-peTrpL in a Tc-dependent complex with MS2-rnTrpL.** Elution samples of MS2-MBP affinity chromatography purification from strain 2011 (pSRKGm-, pRK-MS2-rnTrpL), 10 min after induction of 3×FLAG-peTrpL peptide production with IPTG. The elution samples were separated in Tricine-SDS-PAGE, blotted and Western blot analysis with FLAG-specific antibodies was conducted. Presence of Tc in the washing buffer of the purification procedure is indicated. Elution fractions of a control, mock purification from a strain producing the MS2 RNA aptamer instead of MS2-rnTrpL were also analyzed. Lane C, positive control, purified, FLAG-tagged FLAG-FimV Protein (Rossmann et al., 2019). **B)** Western blot analysis of 20 ng synthetic 3×FLAG-peTrpL. Prestained protein standard bands are indicated (in kDa). The 3×FLAG-peTrpL band is marked with an arrow.

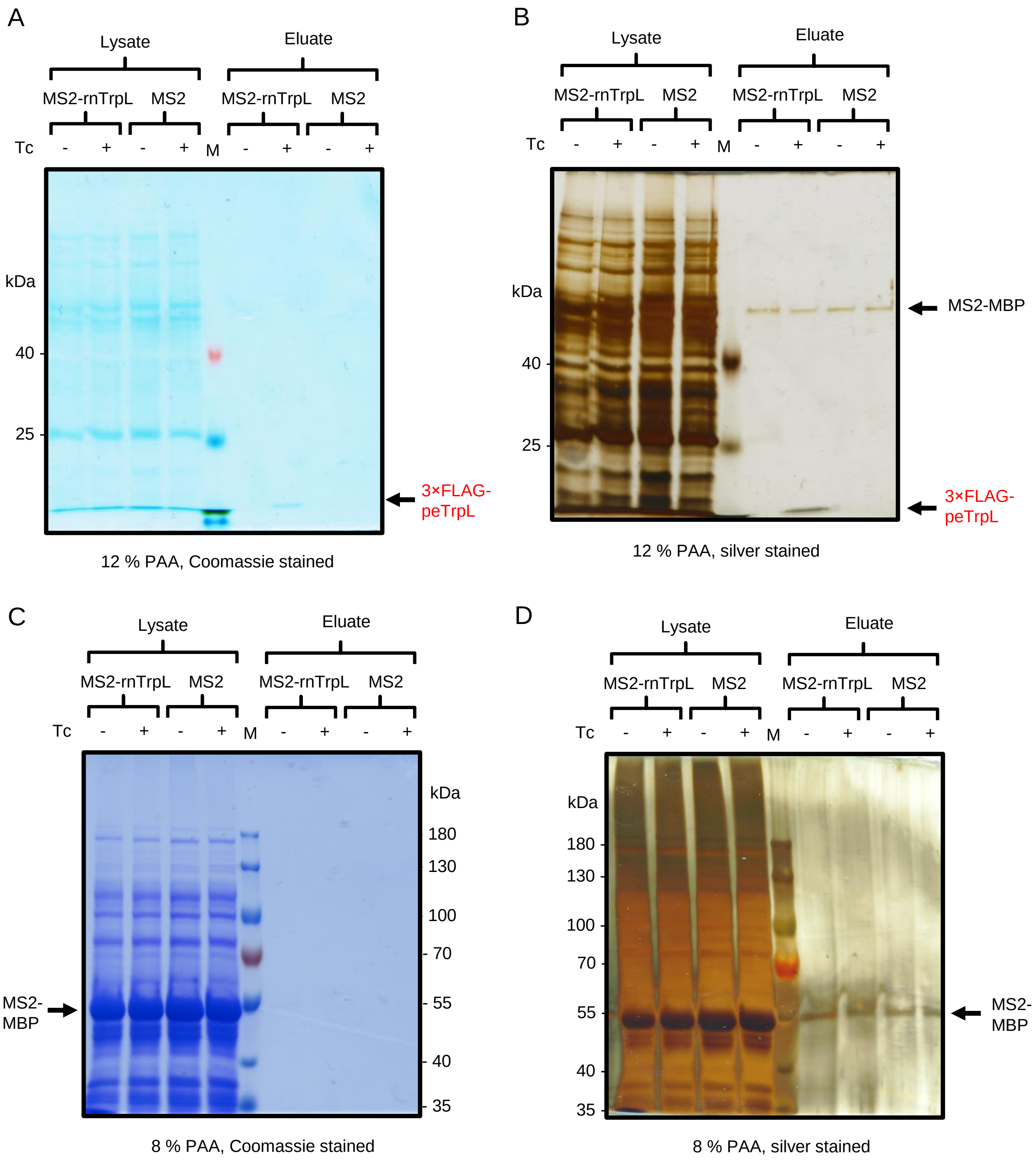


**Figure S6. Glycine-SDS-PAGE analysis of lysates and elution fractions of the MS2-MBP affinity chromatography**. Acrylamide-bisacrylamide (37,5:1) solution was used for the gel preparation. Lysates from cultures grown with 20 µg/ml Tc in the medium were used. The lysates were mixed with MS2-MBP coated amylose beads. MS2-rnTrpL, the tagged sRNA was produced in *S. meliloti* from plasmid pRK-MS2-rnTrpL. MS2, as a negative control, only the MS2-tag was produced from plasmid pRK-MS2. Presence or absence of 2 µg/ml Tc in the lysis and washing buffer of the purification procedure is indicated.

B

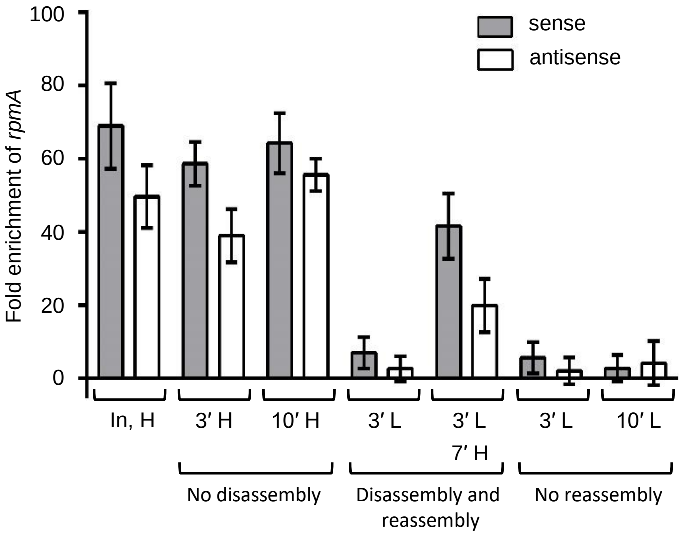


**Figure S7. Tc-dependent reassembly of disassembled complexes containing peTrpL, rnTrpL, and *rplUrpmA* (ARNP).** ARNP containing both the tagged sRNA MS2-rnTrpL and 3×FLAG-peTrpL was purified by MS2-MBP affinity chromatography from strain 2011 (pRK-MS2-rnTrpL, pSRKGm-3×FLAG-peTrpL). The sample containing 3 µg/ml Tc (High-Tc, H) was diluted five-fold and incubated for 3 min at 0.4 µg/ml Tc (Low-Tc, L) for complex disassembly. Then the Tc concentration of the diluted sample was raised to 2 µg/ml and the sample was incubated for 7 min for complex reassembly. Control samples were incubated only at High-Tc (no disassembly) and Low-Tc (no reassembly) conditions, respectively. Then CoIP with 3×FLAG-peTrpL was performed and the *rpmA* level in the CoIP-RNA was monitored by qRT-PCR. The highest amount of *rpmA* mRNA (comparable to the input, In) was coimmunoprecipitated when the ARNP-complex was kept at High-Tc (see bars 3′ H and 10′ H). In contrast, *rpmA* was essentially lost upon incubation in Low-Tc (see bars 3′ L and 10′ L). Importantly, raising the Tc concentration in the diluted sample from Low-Tc to High-Tc led to coimmunoprecipitation of more than the half of the originally co-purified *rpmA*, indicating successful complex reassembly (compare the Input sample to the 3′ L, 7 ′ H sample).

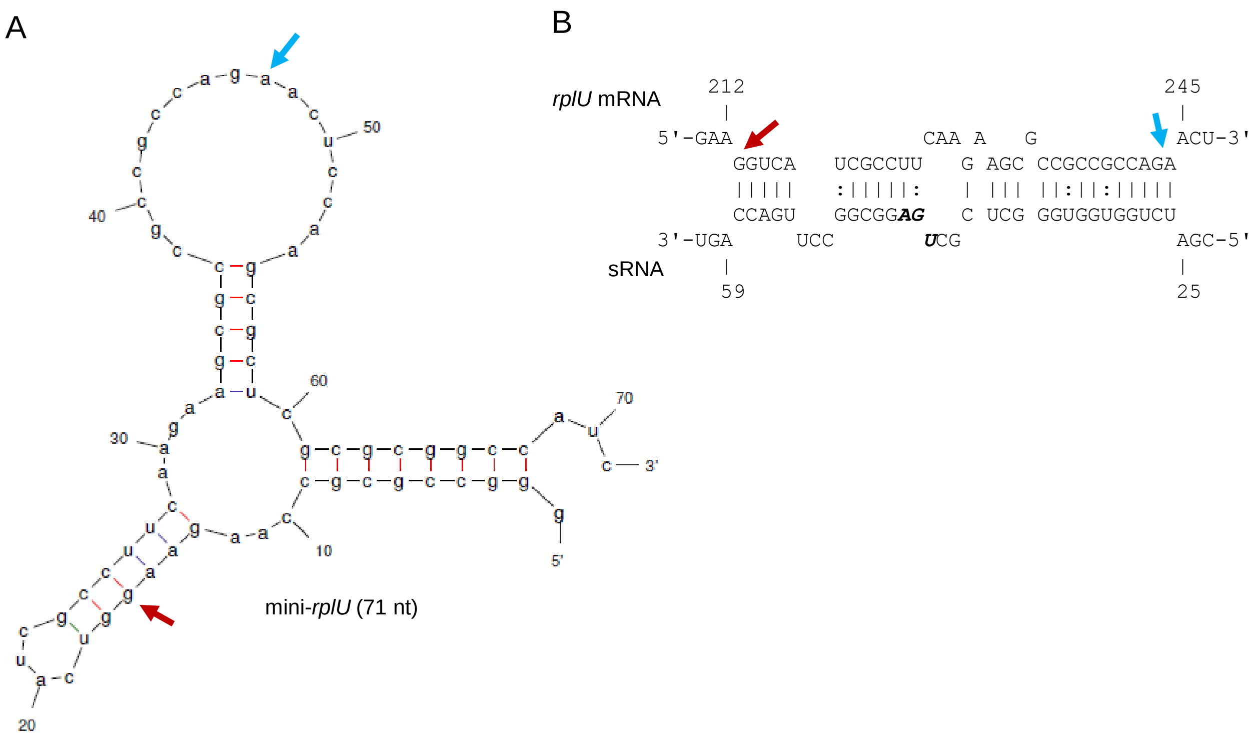


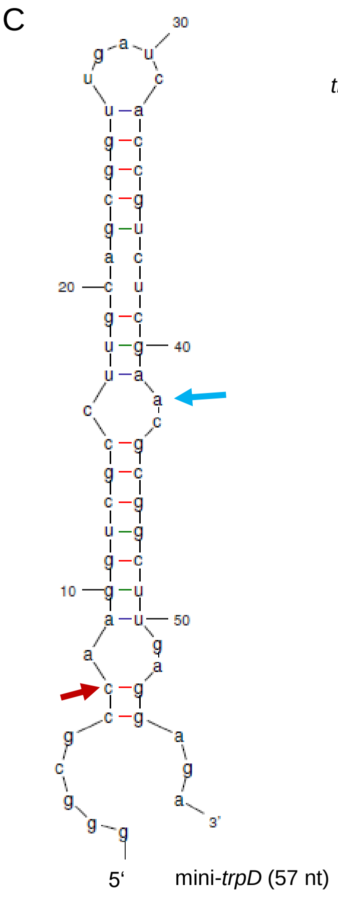


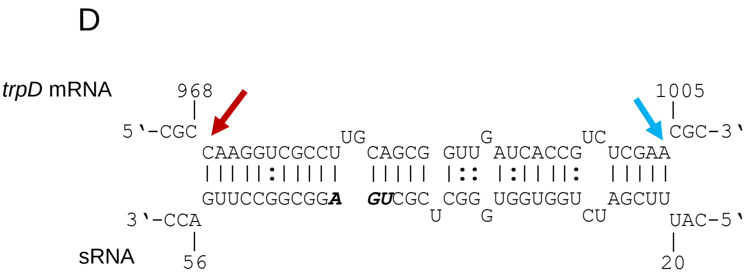


**Fugure S8. Predicted secondary structures of the mini-*rplU* (A) and mini-*trpD* (C) in vitro transcripts and of the interaction of the corresponding mRNAs with rnTrpL (B and D).** Begin and end of the predicted interaction regions are marked with color arrows. The transcripts contain three consecutive guanine at their 5‘-ends, which ensure efficient in vitro transcription by T7 polymerase. According to MFOLD, these guanines do not change the predicted secondary structure of the transcripts.

### Figure S9

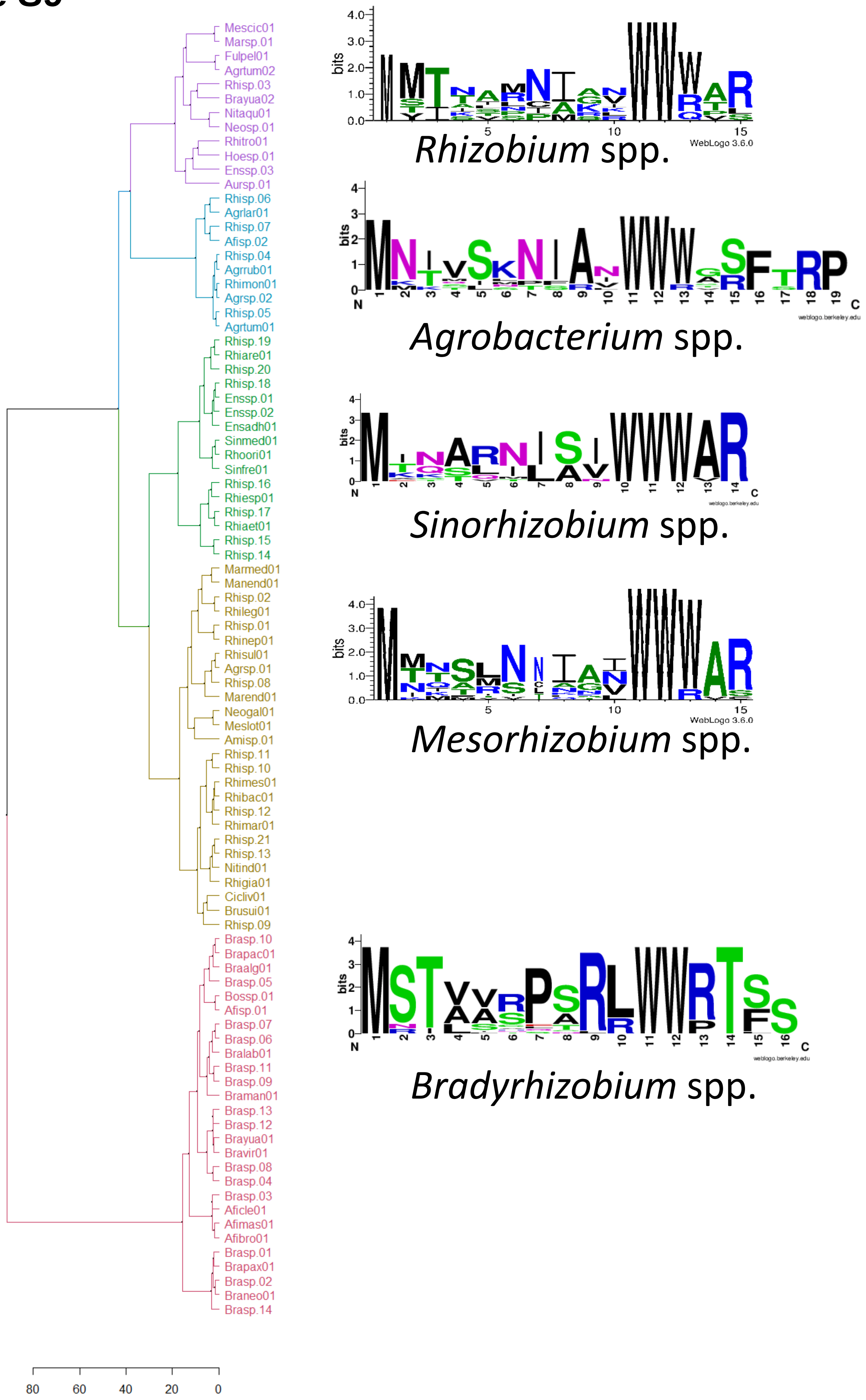
Mescic01
Marsp.01
Fulpel01
Agrtum02
Rhisp.03
Brayua02
Nitaqu01
Neosp.01
Rhitro01
Hoesp.01
Enssp.03
Aursp.01
Rhisp.06
Agrlar01
Rhisp.07
Afisp.02
Rhisp.04
Agrrub01
Rhimon01
Agrsp.02
Rhisp.05
Agrtum01
Rhisp.19
Rhiare01
Rhisp.20
Rhisp.18
Enssp.01
Enssp.02
Ensadh01
Sinmed01
Rhoori01
Sinfre01
Rhisp.16
Rhiesp01
Rhisp.17
Rhiaet01
Rhisp.15
Rhisp.14
Marmed01
Manend01
Rhisp.02
Rhileg01
Rhisp.01
Rhinep01
Rhisul01
Agrsp.01
Rhisp.08
Marend01
Neogal01
Meslot01
Amisp.01
Rhisp.11
Rhisp.10
Rhimes01
Rhibac01
Rhisp.12
Rhimar01
Rhisp.21
Rhisp.13
Nitind01
Rhigia01
Cicliv01
Brusui01
Rhisp.09
Brasp.10
Brapac01
Braalg01
Brasp.05
Bossp.01
Afisp.01
Brasp.07
Brasp.06
Bralab01
Brasp.11
Brasp.09
Braman01
Brasp.13
Brasp.12
Brayua01
Bravir01
Brasp.08
Brasp.04
Brasp.03
Aficle01
Afimas01
Afibro01
Brasp.01
Brapax01
Brasp.02
Braneo01
Brasp.14
80
60
40
20
0
Rhizobium spp.
Agrobacterium spp.
Sinorhizobium spp.
Mesorhizobium spp.
Bradyrhizobium spp.
bits
WebLogo 3.6.0
weblogo.berkeley.edu

**Figure S9. Analysis of putative peTrpL peptides from other Rhizobiales members revealed several groups of conserved leader peptides, generally consistent with taxonomy.** For the complete list of species forming the groups shown by colors, see Table S2. Two major groups can be seen, one with two tryptophans and a consensus MSTvvrPsRL**WW**RTss present in *Bradyrhizobium* and related species (the red, bottom branch of the tree), and a larger group of four variants with slightly varying length and considerably varying sequence; clustering in this group is not very robust. The latter usually have three tryptophans within a loosely conserved context ni(a/s)(n/i/v)**WW**WAR. The comparison of the logos with the functional data in Fig. 12 yields a paradoxical observation that, despite functional conservation in *Sinorhizobium*, *Agrobacterium* and *Bradyrhizobium*, aside of the second tryptophan, the patterns of evolutionary and sequence conservation are markedly different.

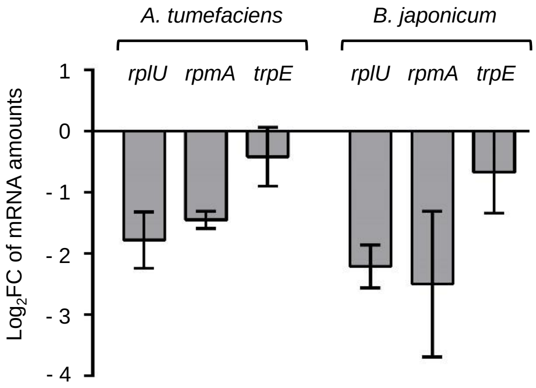


**Figure S10.** qRT-PCR analysis of the indicated mRNAs in *A. tumefaciens* and *B. japonicum*. The *rplUrpmA* mRNA levels were decreased upon overproduction of the respective rnTrpL homologs. Plasmid pSRKTc-Atu-rnTrpL was used to induce LacZ'-Atu-rnTrpL production for 10 min, and mRNA levels after induction were compared to those before induction. Due to the lack of a suitable inducible system for *B. japonicum*, Bja-rnTrpL was overproduced constitutively from the chromosomally integrated, Tc-resistance-conferring plasmid pRJ-Bja-rnTrpL. Shown are means and standard deviations from three independent experiments, each performed in duplicates.
