## Supplementary Methods for "The bacterial leader peptide peTrpL has a conserved function in antibiotic-dependent posttranscriptional regulation of ribosomal genes"

### Melior et al., Supplementary Methods

#### Cultivation of bacteria and conjugation

*E. coli* strains were cultivated in LB supplemented with tetracycline (Tc, 20 µg/ml), gentamycin (Gm, 10 µg/ml) and ampicillin (200 µg/ml) when appropriate. TY was used as growth medium for prototrophic *S. meliloti* 2011 and 2011*ΔtrpL*, and for *A. tumefaciens* NTL4 as described (Melior et al., 2019). Auxotrophic *S. meliloti* 2011*ΔtrpC* strain (Melior et al., 2019) was grown in minimal GMX medium (Schlüter et al., 2010) supplemented with L-tryptophan (Trp) as indicated (Low Trp, 2 µg/ml; High Trp, 20 µg/ml). *B. japonicum* 110spc4 was grown semiaerobically in PSY medium (Mesa et al., 2008). Liquid cultures of Alphaproteobacteria were cultivated semiaerobically (e.g., 30 ml medium in a 50-ml Erlenmayer flask with shaking at 140 r. p. m.) at 30°C to an $OD_{600nm}$ of 0.5, and then processed further. Antibiotics in selective plates or liquid media for Alphaproteobacteria were used at the following concentrations, unless stated otherwise: Tetracycline (20 µg/ml for *S. meliloti* and *A. tumefaciens*; *B. japonicum* was cultivated with 25 µg/ml Tc in liquid and 50 µg/ml Tc on plates), gentamycin (10 µg/ml in liquid cultures and 20 µg/ml in plates), streptomycin (250 µg/ml for *S. meliloti* 2011 and its derivatives), spectinomycin (100 µg/ml for *B. japonicum* 110spc4), chloramphenicol (20 µg/ml in selective plates after plasmid transfer to *B. japonicum*).

The plasmid constructs were transferred by diparental conjugation from *E. coli* S17-1 to *S. meliloti*, *A. tumefaciens* or *B. japonicum* (Simon et al., 1983). Bacteria were mixed, washed in saline and spotted onto a sterile membrane filter, which was placed onto a TY plate without antibiotics. After incubation for at least 4 h (for *S. meliloti* and *A. tumefaciens*) or 3 days (for *B. japonicum*) at 30 °C, serial dilutions were spread on agar plates with selective antibiotics. Constitutively overproducing strains were tested immediately after conjugation and were not passaged or stored.

#### Induction with IPTG

To induce transcription from the *lac* promoter of pSRK-plasmids (Khan et al., 2008), IPTG was added to a final concentration of 1 mM for the indicated periods of time.

#### Short-term exposure of *S. meliloti* to antibiotics

For short-term exposure of bacteria to antibiotics, the following concentrations were used. If the cells contained a Tc-resistance plasmid, 20 µg/ml Tc was used. In cells lacking a plasmid with the appropriate resistance gene, subinhibitory concentrations were used: 1.5 µg/ml Tc, 27 µg/ml Em, 9 µg/ml Cl, 3 µg/ml Rf, 45 µg/ml Km. To ensure that the observed effects were due to the antibiotics and not their solvents, a larger culture was divided into two 30-ml aliquots before the treatment. The first aliquot (the treated culture) was supplemented with 60 µl of an antibiotic stock solution suitable to reach the desired final concentration of the antibiotic. The second aliquot (the control culture) was supplemented with 60 µl of the solvent (ethanol, methanol or water). After the desired exposure time (as indicated; usually 10 min), cells were harvested and processed. To analyze the antibiotic effect, the treated culture was compared to the control culture.

#### Cloning procedures and plasmid characteristics

Plasmid preparation, restriction analysis, purification of DNA fragments from agarose gels, and cloning procedures were performed as described by Sambrook et al. (1989). FastDigest restriction enzymes and Phusion polymerase were used routinely for cloning in *E. coli* DH5alpha. PCR amplicons were first cloned in pJet1.2/blunt (CloneJet PCR Cloning Kit) and the inserts were then subcloned into the conjugative, broad-host range vectors pRK4352 (Mank et al., 2012), pSRKGm or pSRKTc (Khan et al., 2008), which can replicate in *S. meliloti* and *A. tumefaciens*, or in pRJPaph-MCS (Hahn et al., 2016), which was used for chromosomal integration in *B. japonicum*. Alternatively, for very short inserts (e.g. *trpL* ORFs with codons exchanged for synonymous codons, or with mutated codons), complementary oligonucleotides were annealed and the resulting double-strand DNA with suitable single-stranded, cohesive ends, was cloned directly into the desired conjugative plasmids. Insert-containing plasmids were analyzed by Sanger sequencing with plasmid-specific primers (sequencing service by Microsynth Seqlab, Göttingen, Germany) prior to conjugation. All oligonucleotides (primers) are listed in Table S3. They were synthesized by Microsynth, Balgach (Switzerland). The used plasmids are listed in Table S4.

pRK-rnTrpL (Melior et al., 2019) and pRK-MS2-rnTrpL are pRK4352 derivatives that mediate a constitutive transcription of rnTrpL or MS2-tagged rnTrpL (MS2-rnTrpL) in *S. meliloti*. Plasmids pRK-rnTrpL-AU1,2UA, pRK-rnTrpL-CG40,41GC and pRK-rnTrpL-GG46,47CC are pRK-rnTrpL derivatives harboring the indicated mutations in the rnTrpL sequence (Melior et al., 2019). In the rnTrpL-AU1,2UA sequence, the start codon was exchanged for a stop codon, thus inactivating the *trpL* sORF. The CG40,41GC mutation causes an Arg/Ala exchange in peTrpL and weaker base-pairing with *rplU*. The mutation GG46,47CC, which is located downstream of the *trpL* stop codon, also weakens the base-pairing with *rplU.*

Plasmids pSRKTc-rnTrpL and pSRKGm-rnTrpL allow for IPTG-inducible transcription of the recombinant rnTrpL derivative lacZ′-rnTrpL (Melior et al., 2019). In contrast to the leaderless wild type rnTrpL, which starts directly with the ATG of *trpL* (Bae and Crawford, 1990), this rnTrpL variant harbors the *lacZ* 5′-UTR with a ribosome-binding site. Its capability to act in *trans* and to downregulate *trpDC* was shown previously (Melior et al., 2019). Plasmid pSRKGm-peTrpL allows for IPTG-inducible production of peTrpL It contains the sORF *trpL* with several synonymous nucleotide substitutions (to avoid RNA-based effects), but without rare codons (to avoid toxicity; Zahn, 1996). Plasmid pSRKGm-3xFLAG-peTrpL harbors N-terminally fused codons for a triple FLAG-tag. Plasmids pSRKTc-Atu-rnTrpL and pSRKTc-Atu-peTrpL were used for IPTG-induced overproduction of the *A. tumefaciens* rnTrpL and peTrpL homologs. Plasmids pRJ-Bja-rnTrpL and pRJ-Bja-peTrpL were used for constitutive overproduction of *B. japonicum* rnTrpL and peTrpL homologs.

To construct pSRKGm-rplUrpmA′-egfp, primers NdeI-rplU-fw and 5′ -egfp-rpmA-re were used to amplify the *rplUrpmA′* part of the construct, while *egfp* was amplified with primers rpmA-egfp-fw and XbaI-egfp-re. For overlapping PCR, primers NdeI-rplU-fw and XbaI-egfp-re were used, and the PCR product encompassing *rplUrpmA′::egfp* was cloned into pJet1.2/blunt and then subcloned into pSRKGm. IPTG-induced transcription of the bicistronic *rplUrpmA′::egfp* mRNA from pSRKGm-rplUrpmA′-egfp allows for production of L21 and the fusion protein L27′-EGFP (Fig. 4B), which contains only the first three N-terminal aa of L27 fused to the third aa of EGFP. To introduce compensatory mutations into the *rplUrpmA′::egfp* reporter, site-directed mutagenesis of pJet-rplUrpmA′-egfp was performed. Primer pair rplU-G-228-C-FW and rplU-G-228-C-RV was used for one nucleotide exchange, thus re-establishing the base pairing or *rplU* to the nucleotide at position 40 in rnTrpL (re-establishing the influence of rnTrpL-CG40,41GC on *rplUrpmA*; see Fig. 4A). Further, primer pair rplU-CC-222-GG-FW and rplU-CC-222-GG-RV was used for compensatory mutations that re-established the base pairing with rnTrpL-GG46,47CC. The mutated inserts were then subcloned to produce pSRKGm-rplU-CC221,222GG-rpmA’-egfp and pSRKGm-rplU-G228C-rpmA’-egfp.

To construct pSUP-Pas-rpmA-egfp, 190 bp upstream of the putative TSS of the antisense RNA anti-*rplUrpmA* (according to the RNA-seq data, see also Fig. 6C) was amplified using primers EcoRI-asP-rpmA-fw and Pas-rpmA-sSD-rv. The first two nucleotides downstream of the putative TSS were included in the reverse primer, which in addition contained the sequence of a typical bacterial ribosome-binding site and the 5′-sequence of *gfp*. The *gfp* sequence of plasmid pLK64 (McIntosh et al., 2008) was amplified with primers rpmA-sSD-egfp-fw and PstI-egfp-rev, and the two PCR products were used for overlapping PCR with primers EcoRI-asP-rpmA-fw and PstI-egfp-rev. The resulting PCR product was cloned into plasmid pSUP202pol4-exoP, which is a pSUP202pol4 (Fischer et al., 1993) derivative containing 300 nt of the 3′ *exoP* region as a suitable chromosomal integration site (Schlüter et al., 2015). The resulting plasmid pSUP-Pas-rpmA-egfp was used to analyze the Tc-induced transcription from a putative antisense promoter located downstream of *rpmA*. Since this plasmid confers Tc resistance, it was necessary to incubate *S. meliloti* strains with the chromosomally integrated plasmid overnight without Tc (essentially all cells retained the plasmid, as confirmed by qPCR analysis), before adding Tc again to test for induced promoter activity by qRT-PCR analysis of the reporter *egfp* mRNA.

**EGFP fluorescence measurement**

*S. meliloti* 2011*ΔtrpL* strains containing two plasmids (pSRKGm- and pSRKTc- constructs for IPTG-induced expression of *egfp* reporter fusions and the sRNA, respectively), were cultivated in TY with Gm and Tc to an $OD_{600nm}$ of 0.5 and then the production of an EGFP fusion protein and the regulatory sRNA was induced simultaneously with 1 mM IPTG for 20 min. Then, 300 µl of the cultures were transferred to a 96-well microtiter plate and EGFP fluorescence was measured using a Tecan Infinite M200 reader. ODs were also measured using the Tecan reader and used for normalization.

**RNA purification**

To purify total RNA of *S. meliloti* and *A. tumefaciens* for Northern blot hybridization and qRT-PCR analysis, 15 ml culture ($OD_{600}$ = 0.5) was filled into tubes with ice rocks (corresponding to a volume of

15 ml) and pelleted by centrifugation at 6,000 g for 10 min at 4 °C. The pellet was resuspended in 250 µl TRIzol. Cells were lysed in a Laboratory Mixer Mill (Retsch MM200) (4 °C) with glass beads, two times for 15 min, interrupted by incubation at 65 °C for 10 min. After addition of 750 µl TRIzol to the samples, RNA was isolated according to the manufacturer's instructions. To remove residual RNases, the isolated RNA was additionally purified with hot-phenol, phenol:chloroform:isoamylalcohol (25:24:1) and chloroform:isoamylalcohol (24:1), and then ethanol-precipitated. For RNA half-live measurements, 1 ml bacterial culture was added to 2 ml RNAprotect Bacteria Reagent (Qiagen) and RNA was isolated using RNeasy columns (Qiagen). RNA from *B. japonicum* was isolated with hot phenol. The washed and dried RNA was resuspended in 30 µl of ultrapure water. RNA concentration and purity were analyzed by measuring absorbance at 260 nm and 280 nm. 10% polyacrylamide-urea gels and staining with ethidium bromide were used to control the integrity of the isolated RNA. For qRT-PCR RNA, 10 µg samples were digested with 1 µl TURBO-DNase for 30 minutes to remove remaining DNA. PCR with *rpoB*-specific primers was performed for each sample, to check for residual DNA. The DNA-free RNA was then diluted to a concentration of 20 ng/µl for the qRT-PCR analysis (see below).

**Northern Blot hybridization**

Total RNA (10 µg) was denatured in urea-formamide containing loading buffer at 65 °C, placed on ice and loaded on a 1-mm thick, 20 x 20 cm, 10% polyacrylamide-urea gel. Separation by electrophoresis in TBE buffer was performed at 300 V for 4 h. Then, the RNA was transferred to a positively charged nylon membrane for 2 h at 100 mA using a semi-dry blotter. After UV-crosslinking of the RNA to the membrane, the membrane was pre-hybridized for 2 h at 56 °C with a buffer containing 6× SSC, 2.5× Denhardt's solution, 1% SDS and 10 µg/ml salmon sperm DNA. Hybridization was performed with radioactively labeled oligonucleotides (see Table S3) in a solution containing 6× SSC, 1% SDS, 10 µg/ml salmon sperm DNA for at least 6 h at 56 °C. The membranes were washed twice for 2 to 5 min in 0.01% SDS, 5× SSC at room temperature. Signals were detected and analyzed using a BioRad molecular imager and Quantity One (BioRad) software. For quantification, the intensity of the sRNA bands was normalized to the intensity of the 5S rRNA. For re-hybridization, membranes were stripped for 20 min at 96 °C in 0.1% SDS.

**Radioactive labeling of oligonucleotide probes**

5`-labeling of 10 pmol oligonucleotide was performed with 5 U T4 polynucleotide kinase (PNK) and 15 µCi [γ-$^{32}$P]-ATP in a 10 µl reaction mixture containing buffer A provided by the manufacturer. The reaction mixture was incubated for 60 min at 37 °C. After adding 30 µl water, unincorporated nucleotides were removed using MicroSpin G-25 columns.

***In vitro* transcription**

For *in vitro* transcription the MEGAshortscript T7 kit (Ambion) was used. The T7 promoter sequence was either integrated in one of the primers for PCR amplification of the template, or it was present in oligonucleotides that were annealed to obtain double-strand template (see Table S3). When a PCR product was used as a template, it was purified and eluted in ultrapure water. Alternatively, complementary oligonucleotides were annealed to obtain a double-strand template. In a transcription reaction, 500 ng of a purified PCR product or annealed oligonucleotides were used according to the manufacturer's instructions (1x T7-polymerase buffer, 7.5 mM ATP, 7.5 mM CTP, 7.5 mM GTP, 7.5 mM UTP, 25 U T7-enzyme mix) for at least 5 h at 37°C. TURBO-DNase (1 µl) was used to remove the DNA template (1 h at 37°C). The *in vitro* transcript was extracted with acidic phenol, precipitated with ethanol and analysed in a 10% polyacrylamide-urea gel after staining with ethidium bromide.

**Strand-specific, real time, quantitative reverse transcriptase PCR (qRT-PCR)**

For analysis of relative steady-state levels of specific RNAs by real-time RT-PCR (qRT-PCR), the Brilliant III Ultra Fast SYBR® Green QRT-PCR Mastermix (Agilent) was used. Routinely, strand-specific analysis was performed as described by Melior et al. (2019). Specifically, 5 µl Master Mix (supplied), 0.1 µl DTT (100 mM; supplied), 0.5 µl Ribo-Block solution (supplied), 0.4 µl water, 1 µl of the reverse primer (10 pmol/µl), and 2 µl RNA (20 ng/µl) were assembled in a 9-µl reaction mixture. After cDNA synthesis, the samples were incubated for 10 min at 96 °C to inactivate the reverse transcriptase. After cooling to 4 °C, 1 µl of the second primer (10 pmol) was added, and real-time PCR was performed starting with 5 min incubation at 96 °C. Used primers and their efficiencies (determined by PCR of serial two-fold RNA dilutions) are listed in Table S3. A spectrofluorometric thermal cycler (Biorad) was used for the qRT-PCR and the quantification cycle (Cq), was set to a cycle at which the curvature of the amplification is maximal (Bustin et al., 2009). As a reference gene for determination of steady-state mRNA levels, *rpoB* (encodes the β subunit of RNA polymerase) was used (Baumgardt et al., 2016). For half-life determination, it was necessary to use the stable 16S rRNA as a reference molecule. To achieve similar Cq of mRNA and 16S rRNA, 2 µl RNA with a concentration of 0.002 ng/µl was used in a 10-µl reaction for real-time RT-PCR with 16S rRNA specific primers. Cq-values of genes of interest and the reference gene were used in the Pfaffl formula to calculate fold changes of mRNA amounts (Pfaffl, 2001). In the case of outlier (e. g. technical duplicate range > 0.5 Cq), the qRT-PCR analysis was repeated. qPCR product specificity was validated by a melting curve after the qPCR-reaction and by gel electrophoresis, and no-template controls were always included.

For qRT-PCR analysis of total RNA, the analysis of the gene of interest (e.g. *rpmA*) and of the reference gene *rpoB* were performed using portions of the same DNA-free RNA sample, and $\log_2$fold changes of mRNA levels after induction by IPTG and/or exposure to antibiotics were determined. For qRT-PCR analysis of coimmunoprecipitated RNA or RNA co-purified with MS2-rnTrpL, the real-time RT-PCR of the gene of interest was performed using a CoIP-RNA sample, while total RNA of the same culture (harvested prior to cell lysis for CoIP) was used for the *rpoB* real-time RT-PCR. Then, the Pfaffl formula was used to calculate the fold enrichment of specific RNAs by CoIP with 3×FLAG-peTrpL or by affinity chromatography with MS2-MBP, in comparison to the corresponding mock purifications.

**mRNA half-life determination**

Stability of mRNA was determined as described (Melior et al., 2019). For transcription termination, rifampicin was added to a final concentration of 600 µg/ml (stock concentration 60 mg/ml in methanol). Aliquots of 2 ml were withdrawn at time points 0, 2, 4 and 6 min and RNA was isolated. Relative levels of specific mRNAs were determined by qRT-PCR analysis with 16S rRNA as a reference. Linear-log graphs were used for half-lives calculation.

**qPCR**

To determine plasmid DNA levels, qPCR with plasmid-specific primers (Table S3) and Power SYBR® PCR Mastermix were performed. The provided PCR master mix included all components necessary for performing real-time PCR except primers, template, and water, which were added as described for qRT-PCR. The qPCR reaction and quantification were performed as described for the qRT-PCR analysis of total RNA.

**Coimmunoprecipitation of RNA with 3×FLAG-peTrpL and MS2-rnTrpL affinity purification**

*S. meliloti* 2011 strains containing the plasmid pSRKGm-3**×**FLAG-peTrpL, which encodes the triple FLAG-tagged leader peptide (3**×**FLAG-peTrpL) under the control of the *lac* promoter, were used for the coimmunoprecipitation (CoIP) 10 min after addition of IPTG. For the mock CoIP control that was performed in parallel, plasmid pSRKGm-peTrpL was used instead. Cell pellets were resuspended in 5 ml buffer B (20 mM Tris, pH 7.5, 150 mM KCl, 1 mM $MgCl_2$, 1 mM DTT) containing 10 mg/ml lysozyme, 2 µg/ml Tc and 1 tablet of protease inhibitor cocktail (Sigma Aldrich) per 40 ml buffer. Cells were lysed by sonication and 40 µl Anti FLAG® M2 Magnetic Beads (Sigma Aldrich) were added to the cleared lysate. After incubation at 4 °C for 2 h, the beads were collected and split into two portions. One sample was washed 3 times with 500 µl buffer B containing an antibiotic (at the indicated concentration), while the second sample was washed with buffer alone. Protease inhibitors were included in the first two washing steps. Finally, the beads were resuspended in 50 µl buffer B. Coimmunoprecipitated RNA (CoIP-RNA) was purified using TRIzol, without subsequent hot-phenol treatment.

First, amylose beads were non-covalently bound to the MS2 coat protein fused to maltose-binding protein (MS2-MBP). Purification of the fusion protein MS2-MBP from *E. coli* via amylose resin and heparin was performed as previously described (Smirnov et al., 2016). Amylose beads from 300 µl suspension (NEB) were repeatedly washed with buffer C (20 mM Tris, pH 8.0, 150 mM KCl, 1 mM $MgCl_2$, 1 mM DTT) and then incubated with 500 pmol MS2-MBP in 1 ml of the same buffer to prepare the beads for the affinity chromatography. *S. meliloti* 2011 strains containing the plasmid pRK-MS2-rnTrpL, which encodes an rnTrpL derivative tagged at the 5'-end with an MS2 aptamer, were used for co-purification of the MS2-tagged sRNA together with its interaction partners. As a control, a mock purification using bacteria containing plasmid pRK-MS2 was performed. When needed, two-plasmid cultures harboring in addition pSRKGm-3×FLAG-peTrpL were used. In such a case, cells were harvested 10 min after induction of 3×FLAG-peTrpL production. Cells from a 60-ml culture were harvested, resuspended in 600 µl buffer C and sonicated. The cleared lysate (400 µl) was mixed with the MS2-MBP-bound amylose beads and incubated for 2 hours at 4 °C on a tumbler. The beads were split into two portions. One portion was washed three times with 1 ml buffer C containing the appropriate antibiotic as indicated, while the second portion was washed with buffer C alone. The MS2-tagged complex was eluted with 250 µl buffer C containing 12 mM maltose. The RNA and protein content of the elution fractions was analyzed. RNA was purified by hot-phenol and analyzed by qRT-PCR and Northern blot. The bottom fraction of the hot-phenol extracted sample was mixed with 1.5 ml ice cold acetone, the proteins were precipitated over night at -20 °C and washed twice with acetone before dissolving the air dried pellet in water. Proteins were analyzed by Tricine-SDS-PAGE, Western blot analysis and mass spectrometry.
